# The chromatin, topological and regulatory properties of pluripotency-associated poised enhancers are conserved *in vivo*

**DOI:** 10.1101/2021.01.18.427085

**Authors:** Giuliano Crispatzu, Rizwan Rehimi, Tomas Pachano, Tore Bleckwehl, Sara de la Cruz Molina, Cally Xiao, Esther Mahabir-Brenner, Hisham Bazzi, Alvaro Rada-Iglesias

## Abstract

Poised enhancers (PEs) represent a limited and genetically distinct set of distal regulatory elements that control the induction of developmental genes in a hierarchical and non-redundant manner. Before becoming activated in differentiating cells, PEs are already bookmarked in pluripotent cells with unique chromatin and topological features that could contribute to their privileged regulatory properties. However, since PEs were originally identified and subsequently characterized using embryonic stem cells (ESC) as an *in vitro* differentiation system, it is currently unknown whether PEs are functionally conserved *in vivo*. Here, we generate and mine various types of genomic data to show that the chromatin and 3D structural features of PEs are conserved among mouse pluripotent cells both *in vitro* and *in vivo*. We also uncovered that, in mouse pluripotent cells, the interactions between PEs and their bivalent target genes are globally controlled by the combined action of Polycomb, Trithorax and architectural proteins. Moreover, distal regulatory sequences located close to developmental genes and displaying the typical genetic (*i.e*. proximity to CpG islands) and chromatin (*i.e*. high accessibility and H3K27me3 levels) features of PEs are commonly found across vertebrates. These putative PEs show high sequence conservation, preferentially within specific vertebrate clades, with only a small subset being evolutionary conserved across all vertebrates. Lastly, by genetically disrupting evolutionary conserved PEs in mouse and chicken embryos, we demonstrate that these regulatory elements play essential and non-redundant roles during the induction of major developmental genes *in vivo*.

## Introduction

Poised enhancers (PEs) were originally described in ESC^1,2^ as a rather limited set of distal regulatory elements that, before becoming activated upon cellular differentiation, are already bookmarked in pluripotent cells with unique chromatin and topological features. Briefly, in ESC, PEs are already bound by transcription factors and co-activators (e.g. p300), display high chromatin accessibility and are marked with H3K4me1, which are all typical features of active enhancers (**Fig. S1A)**. However, in contrast to active enhancers, PEs are not marked with H3K27ac, but are bound instead by Polycomb Group protein complexes (PcG^3^) and their associated histone modifications (e.g. H3K27me3). Moreover, it was previously reported that PEs can physically interact with their target genes already in ESC^4^ in a PcG-dependent manner^5^. Most importantly, PEs were shown to be essential for the proper induction of their target genes upon differentiation of ESC into Anterior Neural Progenitor Cells (AntNPC)^5^. The previous epigenetic and topological features could explain, at least partly, why PEs are essential for the induction of major developmental genes. More recently, we also showed that the unique epigenetic, topological and regulatory properties of PEs are genetically encoded and critically dependent on the presence of PE-associated orphan CpG islands (oCGI)^6–9^. Altogether, this led us to suggest that the genetic, epigenetic and topological features of PEs could endow anterior neural loci with a permissive regulatory landscape that facilitates the precise and specific induction of PE target genes upon pluripotent cell differentiation^5,9^. However, the previous characterization of PEs was based on the analyses of a few loci in which ESC were used as an *in vitro* differentiation model. Therefore, it is still unclear whether PEs exist and display essential regulatory functions *in vivo*.

## Results

### Poised enhancers (PEs) display their characteristic chromatin signature in the mouse post-implantation epiblast

PEs were previously identified in mESC grown under serum+LIF (S+L) conditions^5^, which only recapitulates part of the pluripotency states that exist both *in vitro* and *in vivo^10,11^*. Therefore, to gather a more complete view of the full repertoire of pluripotency-associated PEs, we generated the necessary data (i.e. ATAC-seq, H3K27ac ChIP-seq, H3K27me3 ChIP-seq) to identify these regulatory elements in 2i mESC (naïve pluripotency) and EpiLC (epiblast-like cells; formative pluripotency) (**Methods**). Next, PEs identified in S+L mESC, 2i mESC and EpiLC were pooled, resulting in 5704 mouse PEs in total (**Fig. 1A**, **Fig. S1A**). A subset of these PEs, which we refer to as PoiAct enhancers (n=727), gets activated in AntNPC as they overlap H3K27ac peaks identified in these cells^5^ (**Fig. 1A**). The previous data sets generated in S+L mESC, 2i mESC and EpiLC were also used to call active (high chromatin accessibiliy/p300 binding, H3K27ac+; n=14752) and primed enhancers (H3K4me1+, H3K27ac-; n=68681) in mouse pluripotent cells (**Methods**) (**Fig. S1A**). Once these pooled enhancer sets were identified, we investigated their epigenetic profiles during early mouse development (i.e. fertilization (E0) to gastrulation (E6.5)). To this purpose, we mined publically available ATAC-seq and H3K27me3 ChIP-seq data sets^12–14^ and also generated relevant data in mouse E6.5 epiblast (i.e. ATAC-seq, H3K27ac ChIP-seq). Regarding H3K27me3 levels at PEs, we found that this histone modification is especially high in sperm^14^, while it progressively accumulates during oocyte development (**Fig. S1B**). Subsequently, H3K27me3 becomes erased upon fertilization and then it progressively increases until it reaches high levels in the post-implantation epiblast (E5.5-6.5) (**Fig. 1B**), thus resembling how this histone modification accumulates at bivalent promoters^14^. Similarly, chromatin accessibility (i.e. measured by ATAC-seq) progressively increases at PEs following fertilization, reaching its highest levels in the post-implantation epiblast (E6.5) (**Fig. 1B**). Therefore, the chromatin signature that PEs display in mESC (*i.e.* high chromatin accessibility and H3K27me3) is also observed *in vivo* in pluripotent cells from the post-implantation epiblast. Importantly, the PEs remain inactive in the post-implantation epiblast, as they display low H3K27ac. In contrast, and in agreement with our previous observations, the PEs became active and, thus, gained H3K27ac at later developmental stages (e.g. E12.5), particularly in the developing brain (**Fig. S1C**). Interestingly, some PEs remain acetylated in postnatal mouse brain tissues (**Fig. S1C**), suggesting that some of these regulatory elements might contribute not only to the induction but also to the maintainance of gene expression.

**Fig. 1:**
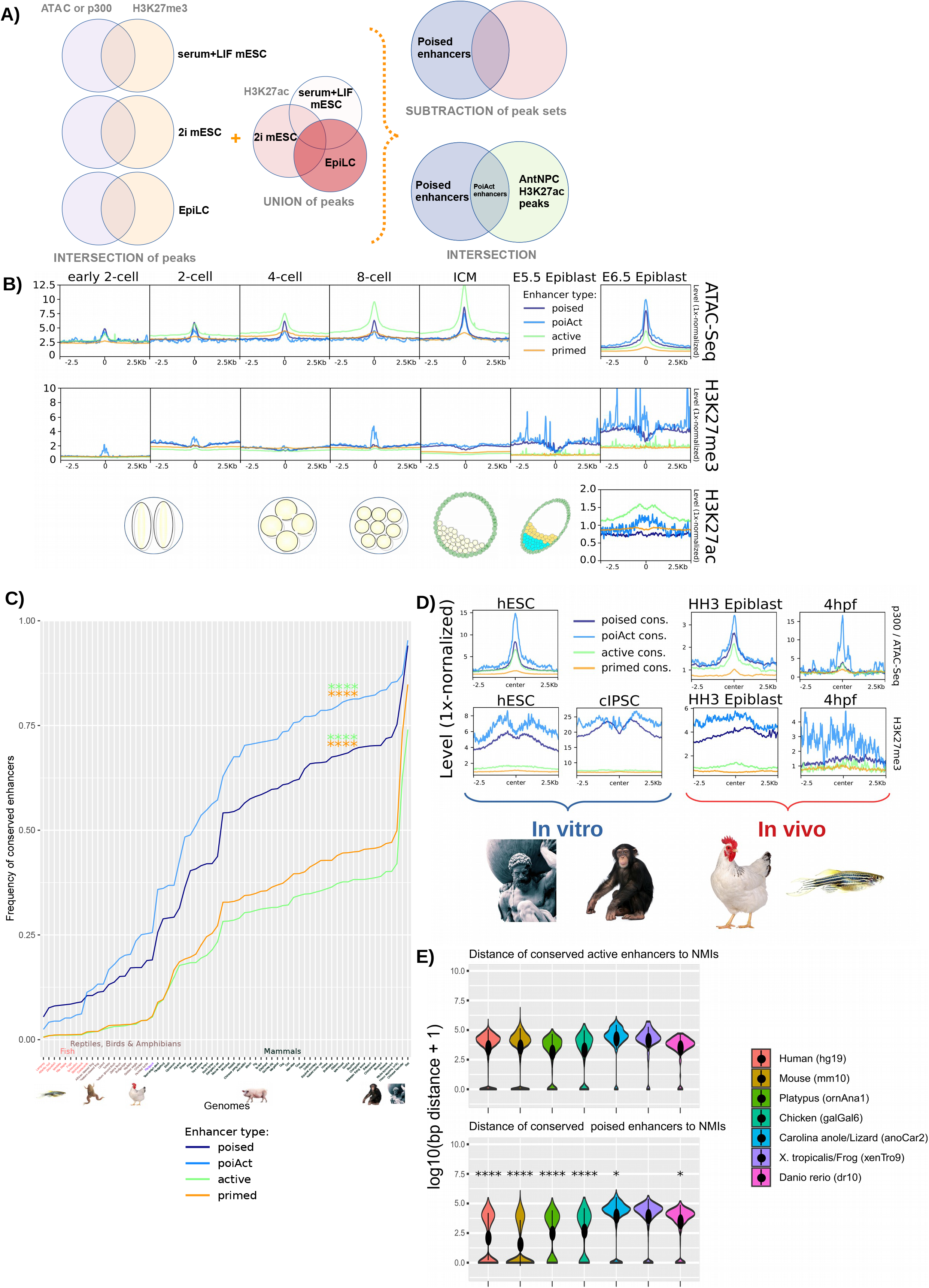
Mouse poised enhancers (PEs) display their characteristic chromatin signature *in vivo* and are highly conserved in mammals. **(A)** ATAC-seq and ChIP-seq data for p300, H3K27me3 and H3K27ac were used to call PEs in three distinct *in vitro* pluripotent states (i.e. 2i ESC, S+L ESC and EpiLC)^*5,79*^. PEs are characterized by high p300/ATAC signals, high H3K27me3 levels and low H3K27ac levels and located at least 5kb apart from a TSS (**Methods**). **(B)** ATAC-seq, H3K27ac and H3K27me3 signals during early mouse embryogenesis^12–14^ (2-cell stage to E6.5 epiblast) are shown around poised, active, primed and PoiAct enhancers identified in *in vitro* pluripotent cells. **(C)** Sequence conservation across 68 vertebrate species (**Table S1**) was measured for poised, active, primed and PoiAct enhancers using a mappability threshold of 0.5. PEs and PoiAct enhancers show significantly higher conservation than active (FC=1.94, p=8.75e-06; FC=2.31, p=9.62e-08 respectively; Wilcoxon test) or primed enhancers (FC=1.68, p=1.092e-05; FC=1.99, p=1.442e-07 respectively; Wilcoxon test). **(D)** ATAC-seq and H3K27me3 signals from human ESC (hESC), chimpanzee iPSC (cIPSC), epiblast from HH3 chicken embryos and 4 hpf zebrafish embryos are shown around poised, active, primed and PoiAct enhancers identified in mouse pluripotent cells and conserved in each of the corresponding vertebrate species. One conserved chicken PoiAct enhancer was removed due to surrounding highly repetitive regions (chr33:3430635–3431044). **(E)** Distance between active (upper panel) or poised (lower panel) mouse enhancers conserved in the indicated vertebrate species and NMIs identified in the same species using Bio-CAP^*6,22*^. The asterisks indicate that PEs are significantly closer to NMIs than active enhancers in all species except the frog (Wilcoxon test). The exact overlap of PEs (compared to active enhancers) with NMIs increases in all these species: in human from 11.63% to 43.4% (p=1.41e-294), in mouse from 11.67% to 56.03% (p<2.2e-16), in platypus from 13.87% to 31.34% (p=6.62e-07), in chicken from 14.56% to 27.8% (p=4.26e-12), in lizard from 4.26% to 10.7% (p=0.0229) and in zebrafish from 5.44% to 10.28% (p=0.019). * p ≤ 0.05, ** p ≤ 0.01, *** p ≤ 0.001, **** p ≤ 0.0001.

### Mouse PEs display high genetic and epigenetic conservation across mammals

Having confirmed that PEs display their characteristic chromatin features *in vivo*, we then assessed their evolutionary conservation among vertebrate genomes (**Methods; Fig. 1C; Fig. S2A-B**), as this can provide preliminary insights into the functional relevance of non-coding sequences^16^. In general, the mouse PEs display high sequence conservation across mammals, which then decreases in non-mammalian amniotes (i.e. birds and reptiles) and is already quite moderate in anamniotes (i.e. amphibians and fish) (**Fig. 1C**, **Fig. S2A-B**). Furthermore, PEs are considerably more conserved than active enhancers across all vertebrates (FC=1.94, p=8.75e-06; Wilcoxon test), probably reflecting the potential involvement of PEs in highly conserved developmental processes (e.g. organogenesis, patterning)^1,5^. Notably, despite their high evolutionary conservation, PEs display a limited overlap with “ultraconserved non-regulatory elements” (uCNEs)^17^ (**Fig. S2C**), which, at least in some cases, act as enhancers of important developmental genes^18–19^. This suggests that, despite their overall high sequence conservation, PEs and uCNEs represent distinct classes of distal regulatory elements.

Having shown that mouse PEs display high genetic conservation, we then wanted to evaluate whether their unique chromatin signature (i.e. high chromatin accessibility/p300 binding, high H3K27me3/PcG binding) was evolutionary conserved. To this end, we mined ATAC-seq and ChIP-seq data previously generated in ESC from different mammals (human^1^, chimp^20^), as well as in early stages of zebrafish development^21^. In addition, we also generated ATAC-seq and H3K27me3 ChIP-seq data for the chicken epiblast (HH3). Next, we analyzed the chromatin features of the mouse PEs that were conserved in each of the previous vertebrate species (**Fig. 1D**). In mammalian ESC and to a lesser extent in the chicken epiblast, PEs showed the expected chromatin signature i.e. high H3K27me3 and ATAC-seq levels, while this was less obvious in zebrafish embryos, especially for H3K27me3. There are probably several reasons contributing to these differences: (i) the conserved PEs in non-vertebrate species might include TF binding sites (indirectly detected by p300/ATAC peaks) but not nearby oCGI, which are responsible of PcG recruitment and H3K27me3 enrichment^9^ (see below); (ii) the lower number of conserved PEs in non-mammalian vertebrates (chicken n=982, zebrafish n=459) can make the ATAC-seq and ChIP-seq profiles to look noisier; (iii) the data for non-mammalian vertebrates was generated *in vivo*, which due to lower cell numbers and higher cellular heterogeneity can also compromise the quality of the ATAC-seq and ChIP-seq profiles. Nevertheless, even in non-mammalian vertebrates (i.e. chicken and zebrafish), the conserved PEs display ATAC-seq and H3K27me3 signals clearly higher than those observed for other enhancer classes.

### PEs are a prevalent feature of vertebrate genomes

We previously found that PEs have a unique and modular genetic composition^5,9^, since compared to other enhancer classes, they are frequently located close to oCGI (i.e. 70%-80% of PEs are located within 3 kb of a computationally defined “weak” CGI^5^ or a biochemically defined CGI^9^, also known as Non-Methylated Islands (NMI)^22^). Our previous work based on the in-depth analyses of a few selected PE loci suggests that the proximity to CGI confers PEs with unique epigenetic features, such as binding by PcG and DNA hypomethylation^5,9,23–25^. Congruently, analysis of whole-genome bisulfite sequencing data generated in mouse pluripotent cells^26,27^ revealed that PEs are globally hypomethylated both *in vitro* and *in vivo* (**Fig. S3A**). Moreover, PEs are bound in mESC by KDM2B (**Fig. S3B**), a protein containing CXXC domains that specifically recognize CGI and that might be responsible, at least partly, of the unique epigenetic properties of the PEs^24,28,29^ (**Fig. S3C**). Given the importance that CGI might have in conferring PEs with their unique chromatin and regulatory features, we then wanted to evaluate whether the proximity between PEs and CGI was evolutionary conserved. However, CGI have been traditionally identified using algorithms originally implemented in mammalian genomes, but that, due to the variability in overall GC and CpG contents, do not perform well when applied to cold-blooded vertebrate genomes. To overcome these limitations, we used data previously generated in seven different vertebrates by Bio-CAP (biotinylated CxxC-affinity purification^6,22^), an assay that enables the unbiased identification of CGI (also referred to as NMI). Next, we measured the distance between those mouse PEs conserved in each vertebrate species and the nearest NMI (**Fig. 1E; Fig. S3D**). Among the seven vertebrate species, the proximity of PEs to NMIs was particularly obvious in mammals (mouse, human, platypus) and, to a lesser extent, in chicken. Similarly, we observed that conserved PEs tended to be considerably closer to NMIs than their active counterparts in both mammals and chicken, but not in anamniotes (i.e. frog & zebrafish) (**Fig. 1E**). Therefore, the weaker H3K27me3 enrichment observed at conserved PEs in zebrafish embryos (**Fig. 1D**) could be explained, at least partly, by the frequent absence of nearby oCGI.

Together with our sequence conservation analyses, the previous results suggest that PEs are prevalent in mammals but rather scarce in other vertebrates, especially in anamniotes. Alternatively, PEs might be abundant in all vertebrates, but sequence conservation, including the proximity to oCGI, might preferentially occur within individual vertebrate clades. To distinguish between these two possibilities, we called PEs *de novo* in human ESC, chicken epiblast and zebrafish embryos using available epigenomic data sets^1,21,30^ and similar criteria to the ones used to identify PEs in mouse cells (**Fig. 1A**). Notably, these *de novo* PEs were abundant in both mammals (human n=4009) and non-mammals (chicken n=7306, zebrafish n=2534) and, especially in zebrafish, they displayed stronger ATAC-seq and H3K27me3 signals than those PEs identified through conservation analyses (**Fig. 2A; Fig. 1D**). Accordingly, while in human ESC and chicken epiblast both *de novo* and conserved PEs were similarly close to NMIs, in the zebrafish embryos the proximity to NMIs was dramatically increased for the *de novo* PEs (**Fig. 2B**). Furthermore, in all the investigated species, the *de novo* PE were strongly associated with developmental genes, particularly those involved in neural crest and neural development (**Fig. 2C; Fig. S3E-F**). Overall, these results indicate that PEs showing similar genetic (i.e. proximity to oCGI) and epigenetic (i.e. high ATAC/p300 and H3K27me3 levels) features are prevalent in all vertebrates, but that a large fraction of them might be specific to each vertebrate class/group. In agreement with this possibility, chicken and zebrafish *de novo* PEs were highly conserved among birds/reptiles and fish, respectively, but not in other vertebrates (**Fig. 2D**), thus resembling the preferential conservation of mouse PE within mammals (**Fig. 1C**). Nevertheless, we also noticed that those *de novo* PE showing high sequence conservation tended to be even closer to NMIs (**Fig. S3G**) and, displayed higher H3K27me3 levels (**Fig. 2A**), which according to our recent work^9^, might endow PEs with particularly privileged regulatory properties.

**Fig. 2:**
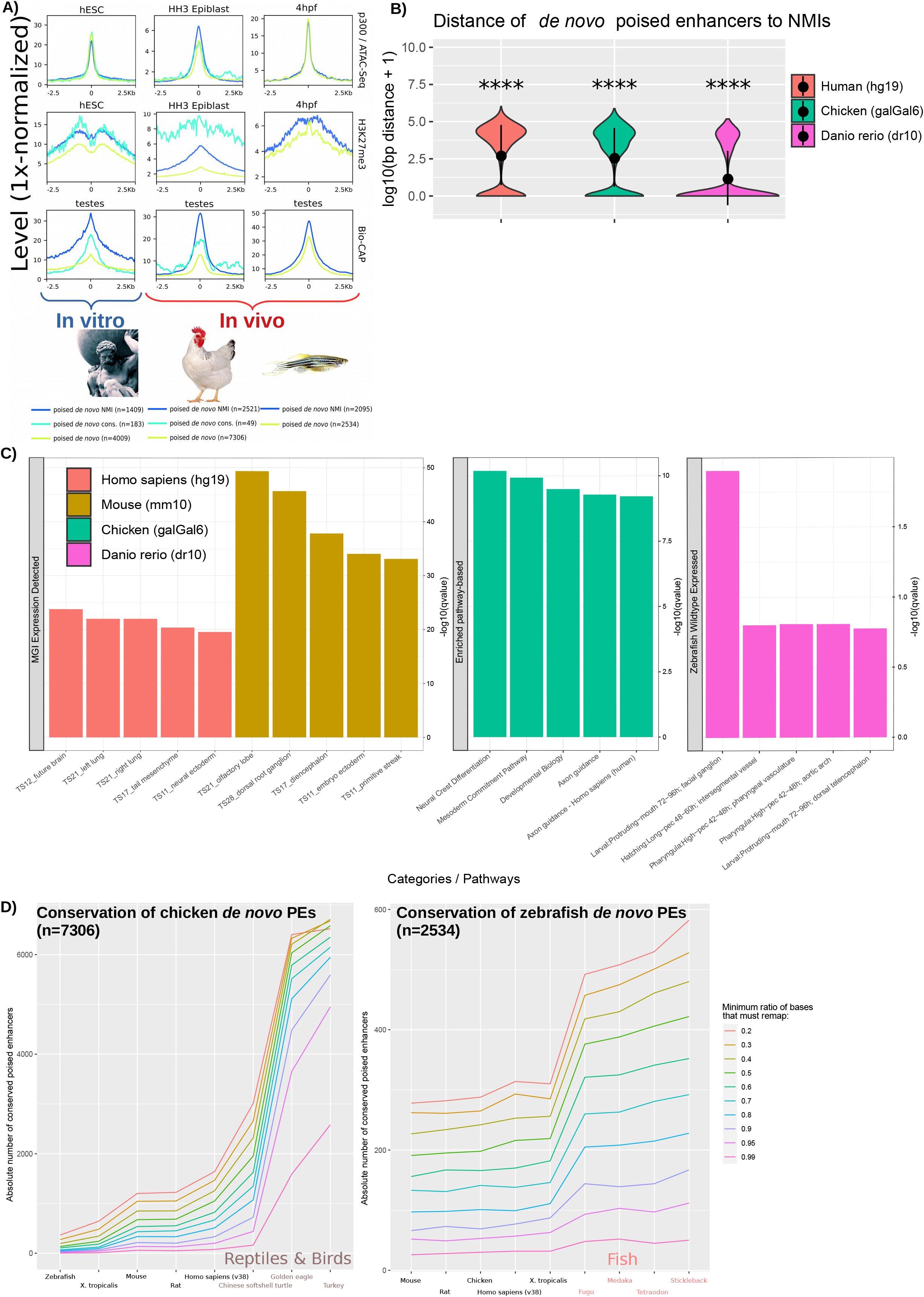
PEs are widespread across higher and lower vertebrates. **(A)** PEs were called *de novo* using available ATAC-Seq, p300 ChIP-seq and H3K27me3 ChIP-Seq data generated in human embryonic stem cells (hESC), HH3 chicken epiblast and zebrafish 4hpf zebrafish embryos (sphere stage; blastulation). ATAC-seq, ChIP-seq and Bio-CAP^*6,22*^ signals are shown for each species around all *de novo* PEs as well as those overlapping NMIs or previously called conserved mouse PEs (except for zebrafish due to low numbers; n=2). **(B)** Distance between *de novo* poised enhancers and NMIs for each of the investigated vertebrate species. The asterisks indicate that *de novo* PEs are significantly closer to NMIs than active mouse enhancers conserved in each species. The exact overlap of *de novo* PEs (compared to active enhancers) with NMIs increases in all species: in human from 11.63% to 33.18% (p=4.15e-56); in chicken from 14.56% to 34.15% (p=3.05e-17) and in zebrafish from 5.44% to 68.77% (p=1.37e-47). **(C)** *De novo* PE sets were annotated using GREAT 3.0^80^, except for the chicken, as this software is not available for this species. The chicken *de novo* PEs were linked to the nearest gene and the resulting gene set was analysed using ConsensusPathDB^81^. **(D)** The sequence conservation of *de novo* PEs identified in chicken (left panel) and zebrafish (right panel) was measured across representative species of the main vertebrate clades using different mappability thresholds (mapping ratios 0.2-0.99; color legend). * p ≤ 0.05, ** p ≤ 0.01, *** p ≤ 0.001, **** p ≤ 0.0001.

### PEs globally interact with bivalent gene promoters in pluripotent cells both in vitro and in vivo

The presence of CGI in the vicinity of PEs provides them with unique topological properties. Namely, PEs can already physically interact with their target genes in ESC, thus preceding their activation in neural progenitors^5,9^. This might confer anterior neural loci with a permissive regulatory topology that facilitates the precise and robust induction of PE target genes. However, this model is based on the detailed analysis of a few endogenous (i.e. *Sox1, Six3, Lmx1b, Lhx5*) or ectopic (i.e. *Gata6*) PE loci using 4C-seq technology^5,9^. In principle, the generality of our previous observations could be addressed using Hi-C technology. However, to detect enhancer-promoter contacts at high resolution, Hi-C typically requires large and cost-prohibitive sequencing depths^31,32^. Therefore, we decided to use a more targeted approach, called HiChIP^33^, that combines Hi-C and ChIP, thus allowing interrogation of DNA loops associated with proteins/histone marks of interest. As both PEs and their target gene promoters are typically enriched in H3K27me3^1,5^, we first generated HiChIPs for H3K27me3 in mESC as biological duplicates. Both replicates were pooled to call loops (p<0.01; n=72265; ranging from 25 Kb-1.81 Mb in loop size; **Methods**). In order to validate the quality of the previous HiChIP data and its usefulness to identify PE-target gene interactions, we first evaluated the four PE loci previously analyzed by 4C-seq and confirmed the reported contacts (**Fig. 3A; Fig. S4A**). Moreover, when considering all the detected HiChIP loops, we found that a large fraction of distal PEs (n=4436; >10kb from a TSS) interacts with at least one locus (55.82%). The majority of these interactions (54.18%) were with gene promoters (**Fig. 3B**), which, importantly, tend to display a bivalent state in mESC (**Fig. 3C**) and are involved in developmental processes (e.g. neural crest and brain development, patterning, morphogenesis) (**Fig. 3D**). Interestingly, we noticed that, in contrast to the frequent long-range/inter-TAD interactions established between PcG-bound genes/domains^3,4,34^, PE-gene contacts preferentially occur within the same TAD (i.e. intra-TAD) (**Fig. 3E**). Moreover, the interactions between PEs and bivalent genes detected by HiChIP were also readily observed in Hi-C data generated in mESC grown under both serum+LIF and 2i conditions^35^ (**Fig. 3F**). This further validates the quality of our HiChIP data and supports that PE-gene contacts are present across different *in vitro* pluripotent states (**Fig. S1A**). Once these pre-formed contacts between PEs and their target genes in mESC were globally confirmed, we also investigated whether this topological feature could be also observed *in vivo* using Hi-C data recently generated in peri-implantation mouse embryos (E3.5-E7.5)^36^. Notably, we observed clear contacts between PEs and bivalent promoters in the mouse ICM as well as in the post-implantation epiblast (**Fig. 3G**). This is in agreement with the overall conservation of the PEs chromatin features *in vivo* (**Fig. 1B**), since such features, namely H3K27me3/PcG, are considered as important mediators of PE-gene interactions^5^.

**Fig. 3:**
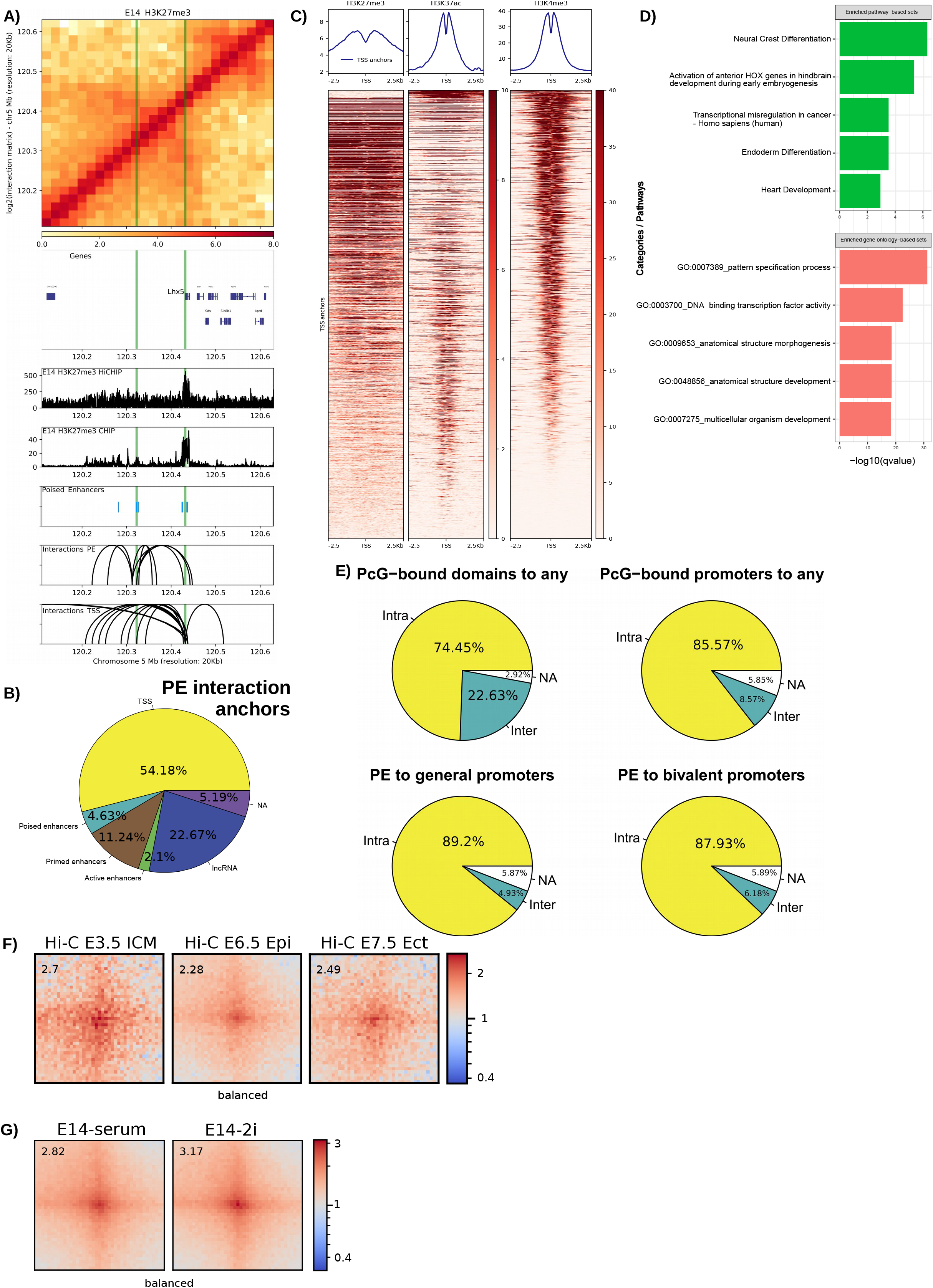
Mouse PEs globally interact with bivalent promoters in pluripotent cells both *in vitro* and *in vivo*. **(A)** H3K27me3 HiChIP and ChIP-seq profiles generated in mESC are shown around the *Lhx5* locus. Both the *Lhx5* TSS and a PE previously shown to control this gene in AntNPC (PE *Lhx5* (−109kb)) are marked with green vertical lines. <u>Upper panel:</u> heatmap of log2 interaction intensities based on H3K27me3 HiChIP data generated in mESC. <u>Two medium panels:</u> ChIP-seq and 1D HiChIP signals for H3K27me3 in mESC. <u>Lower two panels:</u> significant (p<0.01; FitHiChIP^82^) interactions were called using the H3K27me3 HiChIP PETs (paired-end tags) and loops in which one of the anchors overlaps (+/− 10kb) either the PE *Lhx5* (−109kb) or the *Lhx5* TSS are shown. **(B)** Significant (p<0.01; FitHiChIP) interactions were called using the mESC H3K27me3 HiChIP PETs (paired-end tags). Interaction anchors were overlapped with PEs and their interaction partners were hierarchically annotated with the categories shown in the pie chart (**Methods**). **(C)** mESC ChIP-seq profiles for H3K27me3, H3K27ac and H3K4me3 are shown around the TSS interacting with PEs according to the H3K27me3 HiChIP loops identified in mESC (p<0.01; FitHiChIP). **(D)** Pathway (green) and Gene Ontology (red) analysis for the TSS/genes interacting with PEs according to the H3K27me3 HiChIP loops identified in mESC (p<0.01; FitHiChIP) was performed with ConsensusPathDB. **(E)** The interactions (called using mESC H3K27me3 HiChIP data; p<0.01; FitHiChIP) established by PcG domains (**Methods**) and PcG-bound promoters as well as those involving PE-general promoter or PE-bivalent promoter pairs were classified as either intra-TAD (yellow) or inter-TAD (green) depending on whether the two anchors of each loop occur within the same or different TADs, respectively. **(F-G)** Significant (mESC H3K27me3 HiChIP data; p<0.01; FitHiChIP) interactions between distal PEs (>10kb from TSS) and bivalent promoters were visualized as pileup plots using Hi-C data generated *in vitro*^35^ (2i ESC, serum+LIF ESC; **F)** and *in vivo*^36^ ( E3.5 Inner Cell Mass (ICM), E6.5 epiblast, E7.5 ectoderm; **G).** Hi-C pairwise interactions are shown 250kb up- and downstream of each PE and bivalent promoter pair. Hi-C interaction matrices were KR-balanced. The numbers in the upper left corners correspond to “loopiness” values (center intensity normalized to the intensity in the corners).

### The physical communication between PEs and their target genes depends on the combined action of Polycomb, Trithorax and architectural proteins

There is some discrepancy regarding which PcG complexes, PRC1 or PRC2, contribute to the physical communication between PcG-bound loci, including PE-target gene interactions^4,5,37–39^. To investigate whether PRC1 and/or PRC2 are globally involved in the establishment of PE-target gene interactions in mESC we used PRC2 (EED^−/−^ mESC^40^) and PRC1 (RING1a^−/−^RING1b^fl/fl^ mESC^41^ treated with Tamoxifen for 72 hours) null mESC lines (**Fig. S4B**). In contrast to H3K27me3, which is globally lost in PRC1 and PRC2 null mESC^5,42^, H3K4me3 levels are maintained or even increased at gene promoters in general and bivalent ones in particular^43^ (**Fig. S4C**), thus enabling the use of this histone mark to study PE-gene interactions in PcG-null ESC. Therefore, we generated H3K4me3 HiChIP data as biological duplicates in each PcG-null mESC line. Overall, we observed that PRC1 null ESC showed a reduction in mid-range interactions (**Fig. 4A**). Most importantly, the contacts between PEs and bivalent promoters were globally reduced in PRC1 null cells (**Fig. 4B**). In contrast, in PRC2 null cells we observed strongly reduced PE-gene interactions within previously investigated loci^**5**^ (**Fig. 4C**, **Fig. S4D**), but only mild effects at a global level (**Fig. 4B**). The importance of PRC1 for PE-gene interactions was also confirmed using Hi-C data generated in different RING1A/B-depleted mESC lines (RING1a^−/−^RING1b^DEG^ mESC) (**Fig. 4D**). Therefore, our global analyses indicate that PRC1 is required for proper PE-target gene communication in ESC. This is likely to be mediated by canonical PRC1 (cPRC1) through the polymerization and/or phase-separation capacity of its PHC and CBX subunits, respectively^34,44–48^.

**Fig. 4:**
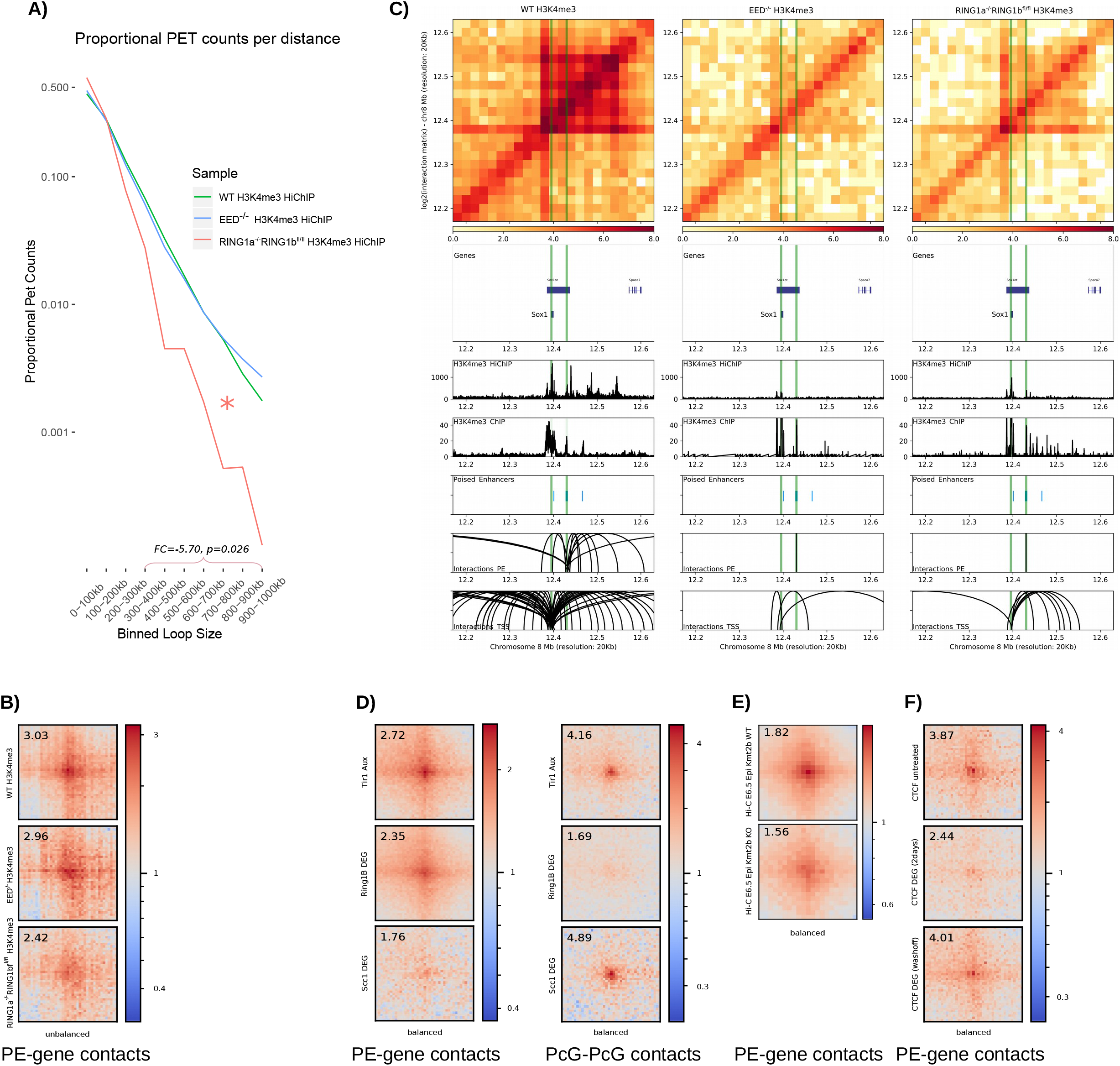
Protein complexes controlling the interactions between PEs and bivalent promoters in mESC. **(A)** Significant (q<0.1; FitHiChIP) H3K4me3 HiChIP interactions identified in WT mESC, *Eed*^−/−^ mESC and Tamoxifen-treated *Ring1a^−/−^Ring1b*^*fl*/*fl*^ mESC were plotted according to their loop size. Mid-range (300kb-1mb) loops are significantly reduced (FC=-5.70, p=0.026; Wilcoxon test) in Tamoxifen-treated *Ring1a*^−/−^*Ring1b*^*fl*/*fl*^ mESC compared to WT. **(B)** Significant (mESC H3K27me3 HiChIP data; p<0.01; FitHiChIP) interactions between distal PEs (>10kb from TSS) and bivalent promoters were visualized as pileup plots using H3K4me3 HiChIP data generated in WT, *Eed*^−/−^ and Tamoxifen-treated *Ring1a*^−/−^*Ring1b*^*fl*/*fl*^ mESC. HiChIP interaction matrices were coverage-normalized. **(C)** H3K4me3 HiChIP in WT, *Eed*^−/−^ mESC and Tamoxifen-treated *Ring1a*^−/−^*Ring1b*^*fl*/*fl*^ mESC around the *Sox1* locus. **(D)** Significant (mESC H3K27me3 HiChIP data; p<0.01; FitHiChIP) interactions between distal PEs (>10kb from TSS) and bivalent promoters (left panel) or between PcG domains (right panel) were visualized as pileup plots using Hi-C data generated in the following mESC lines expressing *Tir1* and treated with Auxin^51^: WT mESC (*Tir1* Aux), *Ring1a*^−/−^ *Ring1b*^*DEG/DEG*^ mESC (i.e. RING1A/B depleted cells; Ring1b Aux) and Rad21/Scc1^DEG/DEG^ mESC (i.e. Cohesin depleted cells; Scc1 Aux). Hi-C interaction matrices were KR-balanced. **(E)** Significant (mESC H3K27me3 HiChIP data; p<0.01; FitHiChIP) interactions between distal PEs (>10kb from TSS) and bivalent promoters were visualized as pileup plots using Hi-C data generated in the epiblast from WT and *Kmt2b/Mll2*^−/−^ E6.5 mouse embryos^50^. Hi-C interaction matrices were KR-balanced. **(F)** Significant (mESC H3K27me3 HiChIP data; p<0.01; FitHiChIP) interactions between distal PEs (>10kb from TSS) and bivalent promoters were visualized as pileup plots using Hi-C data generated in CTCF-Degron mESC lines^*52*^ that were either untreated (CTCF untreated), treated with Auxin for two days (CTCF Aux (2 days)) or treated with Auxin and then washed off (CTCF Aux (washoff)). Interaction matrices were KR-balanced. In (**B**, **D-F**), HiChIP or Hi-C pairwise interactions are shown 250kb up- and downstream of each PE and bivalent promoter pair. The numbers in the upper left corners correspond to “loopiness” values (center intensity normalized to the intensity in the corners).

It has been recently shown that the long-range interactions between PcG-bound genes/domains in mouse pluripotent cells are controlled not only by PRC1 but also by other protein complexes (i.e. Trithorax/MLL2, Cohesin)^49^. Using Hi-C data generated in E6.5 epiblasts from *Kmt2b*^*ko*^ (*Mll2*^*ko*^) mice^50^, we observed that the contacts between PEs and bivalent promoters were also reduced in the *Kmt2b* mutant epiblasts (**Fig. 4E**). We then analyzed Hi-C data generated in a Scc1/Rad21-Degron ESC line^51^ and found that, while Cohesin depletion lead, as previously reported^51^, to increased long-range interactions between PcG-domains, it actually reduced intra-TAD contacts between PE and their bivalent target genes (**Fig. 4D**). Similarly, CTCF depletion^52^ also diminished the interactions between PE and bivalent genes (**Fig. 4F**). Therefore, PE-bivalent gene contacts might be the result of the combined action of two major mechanisms; (i) homotypic chromatin interactions mediated by PcG and Trithorax complexes and (ii) loop extrusion mechanisms dependent on Cohesin and CTCF that favor PE-gene communication within TADs (**Fig. 3E**).

### The physical interactions between PEs and their target genes are maintained once PE get activated

Our previous analyses based on 4C-seq experiments indicated that PE-gene contacts already present in ESC are maintained in AntNPC once the PE and their target genes become active. However, due to the cellular heterogeneity present within AntNPC, PEs and their target genes get activated only in a fraction of cells that can not be specifically interrogated by 4C-seq/Hi-C experiments. Therefore, we conducted H3K27ac HiChIP experiments in E14 AntNPC^53^ and E10.5 mouse brains to specifically and globally evaluate interactions established by PE that became active (i.e. PoiAct enhancers) in these cells. First, we evaluated the PE loci previously analyzed by 4C-seq and confirmed that PE-target gene contacts are maintained once PE become active in both AntNPC and E10.5 brain cells (**Fig. 5A-B; Fig. S5A**). More importantly, we found that distal PE-bivalent gene contacts detected in mESC (**Fig. 3B**) were also frequently observed once PEs became active in AntNPC (27.47% interacting with at least one locus; P=0.05) or the developing mouse brain (42.58-48.51% interacting with at least one locus; P=0.05) (**Fig. 5C**). Moreover, the PE-gene contacts found in mESC were also detected in the E10.5 brain with a high frequency of PoiAct enhancers previously identified in AntNPC (54.06-62.04% interacting with at least one locus; P=0.05) and genes induced during brain development (**Fig. S5B-D**).

**Fig. 5:**
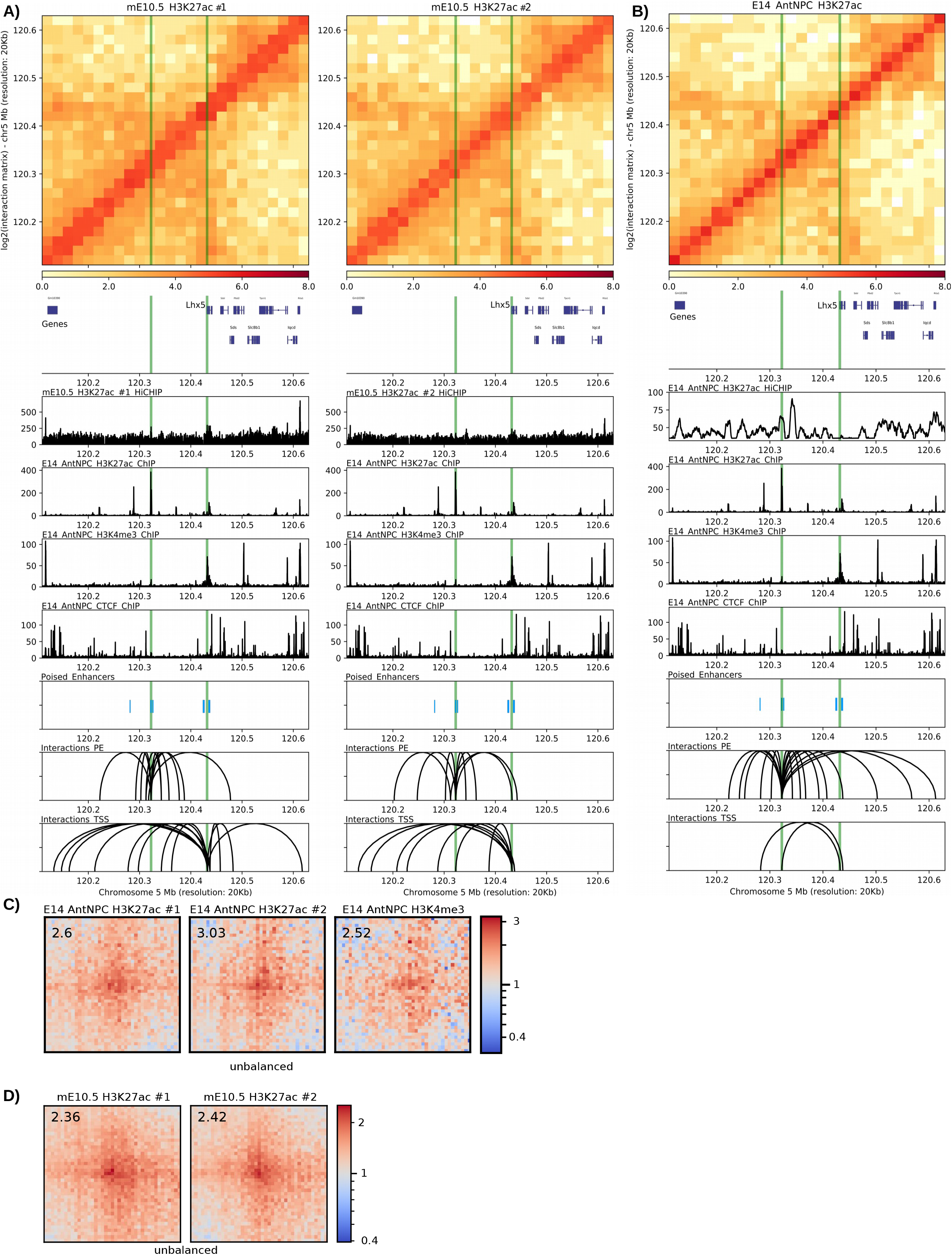
Interactions between PEs and their target genes are maintained once PE become active in either AntNPC or the embryonic mouse brain. **(A-B)** H3K27ac HiChIP and ChIP-seq profiles generated in E10.5 mouse brains (**A**; two biological replicates) and AntNPC (**B**; both replicates merged), respectively, are shown around the *Lhx5* locus. Both the *Lhx5* TSS and the PE *Lhx5* (−109kb) are marked with green vertical lines. <u>Upper panel:</u> heatmap of log2 interaction intensities based on H3K27ac HiChIP data generated in E10.5 mouse brain. <u>Two medium panels:</u> 1D HiChIP signals for H3K27ac in E10.5 mouse brain and H3K27ac, H3K4me3 and CTCF (for potential TAD boundaries) ChIP-seq profiles in AntNPC. <u>Lower two panels:</u> significant (p<0.01 for E10.5 mouse brains; p<0.05 for AntNPC; FitHiChIP) interactions were called using the H3K27ac HiChIP PETs (paired-end tags) and loops in which one of the anchors overlaps (+/− 10kb) either the PE *Lhx5* (−109kb) or the *Lhx5* TSS are shown. **(C-D)** Significant (mESC H3K27me3 HiChIP data; p<0.01; FitHiChIP) interactions between distal PEs (>10kb from TSS) and ESC bivalent promoters were visualized as pileup plots using H3K27ac and H3K4me3 HiChIP data generated in AntNPC (**C**) or H3K27ac HiChIP data generated in E10.5 mouse brain (**D**). HiChIP interaction matrices were coverage-normalized. HiChIP pairwise interactions are shown 250kb up- and downstream of each PE and bivalent promoter pair. The numbers in the upper left corners correspond to “loopiness” values (center intensity normalized to the intensity in the corners).

### Highly conserved PEs are necessary for the induction of major brain developmental genes in vertebrate embryos

The previous analyses show that the main genetic (e.g. proximity to CGI), chromatin (e.g. high H3K27me3/PcG levels) and topological (e.g. pre-formed contacts with target genes) features of PEs are evolutionary conserved and detectable *in vivo*. These observations support, albeit in a correlative manner, the functional relevance of PEs during early vertebrate embryogenesis, particularly during brain development. The functional relevance of PEs was previously demonstrated by deleting candidate PEs in mESC (*i.e. PE Lhx5(−109kb), PE Six3(−133kb), PE Sox1(+35kb), PE Wnt8b(+21kb))*, which severely compromised the induction of major brain developmental genes (*i.e. Lhx5, Six3, Sox1, Wnt8b*) upon differentiation of ESC into AntNPCs^5^. However, it is currently unknown whether PEs also have essential and non-redundant regulatory functions *in vivo* or whether, alternatively, these privileged regulatory properties might represent an *in vitro* “artifact” due to the reduced robustness of *in vitro* differentiation systems. To start addressing this important question, we first used CRISPR/Cas9 technology to generate mouse embryos in which we deleted the *PE Lhx5(−109kb),* one of the PEs that we previously characterized *in vitro* (**Fig. 6A-C; Fig. S6A-B**). Remarkably, the expression of *Lhx5* was strongly reduced in the forebrain of E8.5 and E9.5 *PE Lhx5(−109kb)*^−/−^ mouse embryos in comparison to their WT isogenic controls (**Fig. 6C**). Next, since the mouse *PE Lhx5(−109kb)*^−/−^ has a high genetic and epigenetic conservation across vertebrates, we decided to generate targeted deletions of its homologous sequence in the developing brain of chicken embryos using CRISPR/Cas9^54^ (**Fig. 6C-E**). Briefly, the forebrain of HH9 chicken embryos was unilaterally electroporated with vectors expressing Cas9 and gRNAs flanking the *PE Lhx5* conserved sequence (**Fig. 6D**; **Fig. S6A,C)**. Subsequently, the expression of *Lhx5* was evaluated by *in situ* hybridization (ISH) in HH14 chicken embryos (~ to E11.5 in mice). Notably, the forebrain expression of *Lhx5* was strongly and specifically reduced in the electroporated side (**Fig. 6F**). Furthermore, the specificity of these results was further supported by experiments in which the electroporation of chicken embryos with Cas9 and scrambled gRNAs did not affect *Lhx5* expression (**Fig. 6F**). To further evaluate the *in vivo* functional relevance of evolutionary conserved PEs, we similarly disrupted another two PEs (*PE Six3(−133kb), PE Sox1(+35kb)*) that were previously characterized in mESC^5^ and that were also genetically and epigenetically conserved in the chicken genome (**Fig. S6C-E**). We again observed a strong reduction in the expression of *Six3* and *Sox1* in the electroporated side of embryos targeted with both Cas9 and gRNAs flanking the PEs, but not when Cas9 was electroporated with scrambled gRNAs (**Fig. 6F**). Furthermore, in the case of the *PE Six3(−133kb)* chicken homolog, its disruption resulted in a smaller and malformed eye, in agreement with the strong expression and conserved function of *Six3* during eye development^*55*^ (**Fig. 6F**). Overall, these results demonstrate that the regulatory function of PEs is essential and conserved *in vivo*.

**Fig. 6:**
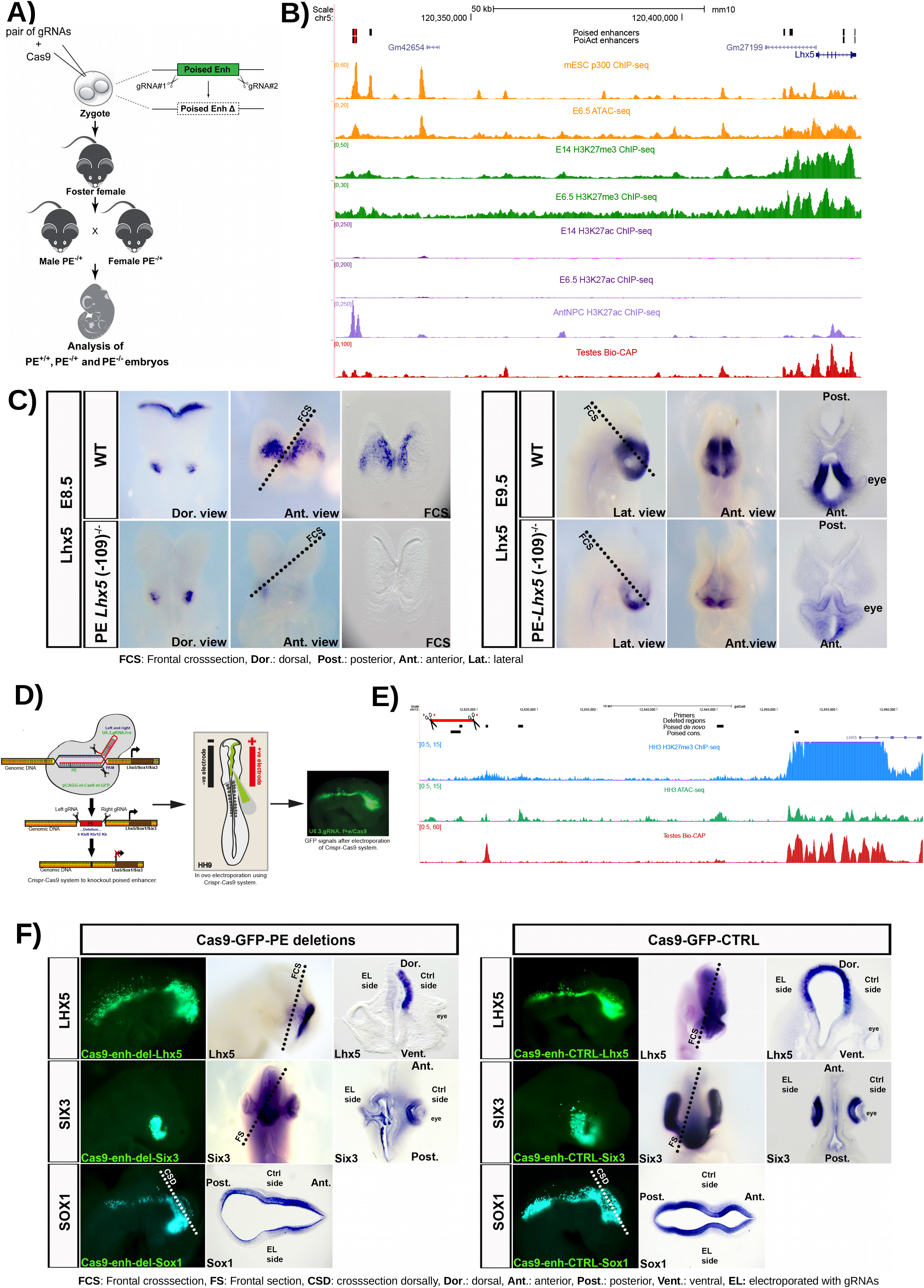
Functional relevance of PEs during *in vivo* brain development. **(A)** Graphical overview of the CRISPR/Cas9 experimental strategy used to generate mouse embryos with homozygous deletions of the PE *Lhx5* (−109kb). **(B)** ATAC-seq and ChIP-seq (p300, H3K27me3 and H3K27ac) profiles generated in mESC and E6.5 mouse epiblast are shown around the *Lhx5* locus. In addition, the Bio-CAP profiles generated in mouse testes is shown to illustrate the location of CGI/NMI in the mouse genome, as well as H3K27ac ChIP-seq profiles generated in AntNPC to illustrate the activation of PE *Lhx5* (−109kb) in AntNPC. The genomic region deleted in mouse embryos that includes the PE *Lhx5* (−109kb) is highlighted in red. **(C)** RNA *in situ* hybridizations were performed to visualize *Lhx5* expression in WT and PE *Lhx5* (−109kb)^−/−^ mouse embryos at embryonic stages E8.5 (top rows) and E9.5 (bottom rows). **(D)** Graphical overview of the CRISPR/Cas9-based approach used to delete three genetically conserved PEs located within the *Lhx5*, *Sox1* and *Six3* loci (left panel). Schematic diagram of *in ovo* electroporation technique in chick embryo at HH9. Mosaic knock-out chicken embryos were generated by unilateral co-electroporation of CAGG>nls-Cas9-nls-GFP and U6.3>PE *Lhx5*/*Sox1*/*Six3*gRNAf+e vectors into the brain region (medium panel). GFP expression in brain and neural tube of an embryo electroporated with the Cas9 and gRNA-GFP vectors (right panel). **(E)** ATAC-seq and H3K27me3 ChIP-seq profiles generated in HH3 chicken epiblast are shown around the *Lhx5* locus. In addition, Bio-CAP profiles generated in chicken testes are also shown to illustrate the location of CGI/NMI in the chicken genome. The genomic region deleted in chicken embryos that includes the mouse PE *Lhx5* (−109kb) conserved in chicken is highlighted in red. **(F)** RNA *in situ* hybridizations were performed to visualize *Lhx5, Six3 or Sox1* expression in HH9 chicken embryos that were unilaterally electroporated with (i) <u>Left panels:</u> Cas9-GFP and gRNAs flanking the conserved PEs associated with *Lhx5, Six3* and *Sox1* (Cas9-GFP-PE deletions); (ii) <u>Right panels</u>: Cas9-GFP and scrambled gRNA (Cas9-GFP-CTRL).

## Discussion

PEs were originally identified, mechanistically dissected and functionally characterized in ESC^1,5^. Our previous work suggested that PE could play essential roles during the induction of major developmental genes once pluripotent cells start differentiating, particularly towards the neural lineage^5^. However, it was still unclear whether PEs actually existed and were functionally relevant *in vivo.* Here we addressed this important question by mining various genomic and epigenomic data sets, which enabled us to conclusively show that PEs not only display their characteristic chromatin signature *in vivo*, but also that they are a prevalent feature of vertebrate genomes. Interestingly, we found that PEs tend to be highly conserved within specific vertebrate groups (e.g. mouse PEs are highly conserved across mammals; chicken PEs are highly conserved in birds and reptiles), while only a relatively small subset of PEs is conserved across all vertebrates. Nevertheless, in all the investigated vertebrate species, PEs were located close to orphan CGI/NMI^6,22^ and linked to major developmental genes. We recently showed that orphan CGI are an essential component of PEs that, together with TAD boundaries, enables them to precisely and specifically control the expression of developmental genes with CpG-rich promoters^9^. Moreover, we also showed that the main regulatory function of these orphan CGI is to serve as tethering elements that bring PEs and their CpG-rich target genes into physical proximity^9^. Therefore, we propose that the association of distal enhancers with CGI might represent an ancestral regulatory mechanism in vertebrate genomes that enables the precise and specific induction of major developmental genes within large regulatory domains^56–58^. Although CGI are typically considered a specific feature of vertebrate genomes, sequences with similar tethering functions might also exist in invertebrates, where they can also be important for the long-range induction of major developmental genes^59–62^.

As mentioned above, the orphan CGI associated with PEs act as tethering elements physically linking these distal regulatory elements with their target genes. Mechanistically, in undifferentiated ESC, this tethering function seems to be mediated by PcG complexes recruited to the CGI present both at PEs and their target gene promoters^5,9,23,63^. Here we show that these PE-gene contacts are globally dependent on PRC1, while PRC2 seems to preferentially contribute to the interactions occurring within specific loci^5^. These results are in agreement with previous reports indicating that long-range interactions between PcG-loci are mediated by cPRC1 subunits^34,44–48,64^, with PRC2 having a lesser and indirect contribution through its capacity to recruit cPRC1 to its genomic targets^65^. CGI can serve as recruitment platforms for other proteins containing CXXC domains (e.g. TET1, CFP1, MLL2/KMT2B), which are frequently part of important chromatin regulatory complexes (e.g. Trithorax (TrxG))^66–68^. It was recently shown that MLL2/KMT2B, an important component of TrxG complexes, can also contribute to the 3D chromatin organization of bivalent genes in pluripotent cells^50,69^. Interestingly, here we found that MLL2/KMT2B also facilitates the interaction between PE and their bivalent target genes. Therefore, PcG and TrxG complexes might cooperate rather than antagonize each other in pluripotent cells in order to mediate homotypic chromatin interactions within PE loci that facilitate future gene induction^38,47,70,71^. In addition to PcG and TrxG complexes, 3D chromatin organization is largely dependent on the combined effects of Cohesin and CTCF, which are necessary for the formation of TADs and other large regulatory domains through a loop extrusion mechanism^52,72–74^. Although PEs are not directly bound by either Cohesin or CTCF in ESC^5^, we found that PE-gene contacts were diminished when either of these two architectural proteins were degraded. Therefore, loop extrusion might also facilitate the physical interactions between PEs and genes located within the same TAD. This is contrast to the role of Cohesin as a negative regulator of inter-TAD interactions between PcG-bound genes^51^.

Transcriptional and phenotypic robustness during development is believed to require complex regulatory landscapes whereby multiple enhancers redundantly control the expression of major cell identity genes^75–77^. In contrast, using ESC as an *in vitro* differentiation system, we previously showed that PEs can control the induction of genes involved in early brain development in a hierarchical and non-redundant manner^5^. Importantly, using both mouse and chicken embryos as experimental models, we have now confirmed the essential role of PEs for the proper induction of developmental genes during vertebrate embryogenesis. We propose that the privileged regulatory properties of PEs depend, at least partly, on nearby oCGI, which confer these regulatory elements with unique chromatin and topological features^9^. Lastly, it is worth highlighting that PEs are preferentially, but not exclusively associated with neural genes (**Fig. 2C**). Moreover, not all PEs might acquire their unique chromatin and topological features already in pluripotent cells and this might occur later in lineage-restricted multipotent progenitors^78^. Therefore, future work should elucidate the full repertoire of PEs and interrogate their function in different somatic lineages and spatiotemporal contexts.

## Acknowledgements

We thank the Rada-Iglesias lab members for insightful comments and critical reading of the manuscript, Elisabeth Kirst and Janine Altmüller (Cologne Center for Genomics; University of Cologne (UoC)) for technical assistance with next-generation sequencing, and the Regional Computing Center of the UoC (RRZK) for providing computing time on the DFG-funded High-Performance Computing (HPC) system CHEOPS, as well as support. We acknowledge and appreciate the CECAD *in vivo* Research Facility (Branko Zevnik) for the generation and maintenance of the deleted PE *Lhx5* mouse line. We also thank Anton Wutz and Miguel Vidal for generously providing us with the *EED*−/− and *RING1a*−/−*RING1b*^*fl*/*fl*^ mESC lines. Work in the Rada-Iglesias laboratory was supported by CMMC intramural funding (Germany), the German Research Foundation (DFG) (Research Grant RA 2547/1-3), *“Programa STAR-Santander Universidades, Campus Cantabria Internacional de la convocatoria CEI 2015 de Campus de Excelencia Internacional”* (Spain), the Spanish Ministry of Science, Innovation and Universities (Research Grant PGC2018-095301-B-I00) and the European Research Council (ERC CoG*“PoisedLogic“*). Giuliano Crispatzu is supported by funding within the CRU329 (DFG 386793560). Rizwan Rehimi is supported by funding within the SFB829 (DFG 73111208).

## Contributions

GC, RRR, ARI conducted most research and prepared the manuscript. GC conducted bioinformatical analysis and HiChIPs. RRR deleted PEs in HH9 chicken, did *in situ*s and HH3 chicken ChIP- / ATAC-seqs. CX and HB deleted PEs in E8.5 and E9.5 mice. HB and RRR did *in situ* in mice deletions. GC and TP did PcG-null ChIPs and molecular evaluations. TB cultured 2i cells and differentiated EpiLC with subsequent ChIP-seqs. SCM technically assisted with initial HiChIP experiments. EMB ensured *in vivo* mice experiments.

## Methods

### Cell Culture

10cm plates were coated with 0.1% gelatin generally overnight. Cells (E14 WT, *EED*−/−^40^ and *RING1a*−/−*RING1b*^*fl*/fl41^ mESC) were thawed and resuspended in 10ml of medium (serum + LIF). Standard serum + LIF medium contained 500 mL Knock-out DMEM (Gibco 10829-018), 95 mL of filtered ES FBS (Gibco 16141-061), 5.9 mL of antibiotics (*Hyclone SV30079.01*), 5.9 mL Glutamax (Gibco 35050-038), 5.9 mL MEM NEAA (Gibco 11140-035), 4.7 mL titrated LIF (Miltenyi Biotec 130-095-777), and 1.3 mL Beta-mercaptoethanol 55 mM (Gibco 21985-023). Cells were split once (1/3 or 1/4) every two days. *Ring1a*−/−*Ring1b*^*fl*/fl^ mESC were treated with Tamoxifen (OHT; 1mM) for 72h right after one passage and RING1B loss was confirmed by PCR genotyping and Western Blot.

### AntNPC differentiation

ESC were differentiated into AntNPC as previously described^5^ with slight modifications. Briefly, E14 WT ESC grown in serum + LIF were trypsinized with 2mL TrypLE Express (Life technologies). Cells were centrifugated (1000 RPM) and then resuspended in 5 mL of N2B27 with 0.1% of BSA (Life Technologies) and 0.1% of bFgf (PeproTech, 100-18B) to get a single cell suspension. Next, cells were counted with the BioRad TC20 cell counter and 15000 cells / cm^2^ were plated in 10 mL of N2B27 with 0.1% of BSA and 10 ng/mL of bFgf. Cells were differentiated for five days and media was changed every day without any PBS washings in between: Day1 (10 mL of N2B27 with 0.004% of BSA and 10 ng/mL bFgf).; Day2 (10 mL of N2B27 with 0.004% of BSA, 10 ng/mL bFgf and 5 uM Xav939/Wnt inhibitor (Sigma-Aldrich, x3004-5mg)); Day3 (10 mL of N2B27 with 0.004% of BSA and 5 uM Xav939/Wnt inhibitor); Day4 (10 mL of N2B27 with 0.004% of BSA and 5 uM Xav939/Wnt inhibitor). On Day 5, cells were washed 1-2 with PBS, and collected for downstream analyses.Differentiation was assessed using RT-qPCR on a Light Cycler 480II comparing AntNPC d5 vs. E14 WT d0 relative gene expression levels (using the *2*∆Ct method) with house keeping gene (*Eef1a1*), pluripotency markers (*Pou5f1*/*Oct4* and *Nanog*), mesoderm marker (*T*) and ectoderm markers (*Six3* and *Lhx5*). Standard deviations were calculated from technical triplicate reactions and were represented as error bars.

### ATAC-seq

Embryonic tissues were extracted and resuspended into single cells. ATAC-seq was essentially conducted following the protocol from Buenrostro et al.^*83*^. In short, single cells were centrifugated at 5000 g for 5 minutes and supernatant was removed. Cells were then lysed with 100 μl cold lysisl cold lysis buffer (10 mM Tris-HCl, pH 7.4, 10 mM NaCl, 3 mM MgCl_2_, 0,1% IGEPAL CA-630) supplemented with 4 μl cold lysisl of protease inhibitor (1 tablet/2 mL conc.) for at least 15 minutes on ice. Immediately after lysis, nuclei were centrifugated at 6000 g for 10 minutes at 4°C. The resulting pellet was resuspended in transposase reaction mix (25μl cold lysisl 2x TD buffer, 10μl cold lysisl transposase (Illumina), 15μl cold lysisl nuclease-free H_2_O) and incubated for 30 minutes at 37°C. Finally, the sample was purified using Qiagen MinElute PCR purification kit according to the manufacturer’s protocol.

### ChIP

Cells were crosslinked in 1% of formaldehyde for 10 minutes (rotating at RT) with subsequent quenching by glycine (0.125M; rotating at RT). Cells were washed twice with PBS and 25x Roche protease inhibitor, then flash frozen in liquid nitrogen and stored at −80°C. Afterwards cells were thawed for ~30 minutes and lysed in 50 mM Hepes, 140 mM NaCl, 1 mM EDTA, 10% glycerol, 0.5% NP-40 and 0.25% TX-100 together with protease inhibitor (Lysis Buffer 1) for 10 minutes at 4°C while rotating. After centrifugation (5 minutes, 2000rcf, 4°C), the supernatant was discarded and the pellet resuspended in 10 mM Tris, 200 mM NaCl, 1 mM EDTA, 0.5 mM EGTA, protease inhibitor (Lysis Buffer 2) and lysed for 10 minutes at 4°C while rotating. After centrifugation (5 minutes, 2000rcf, 4°C), the supernatant was discarded and the pellet was resuspended in 10 mM Tris, 100 mM NaCl, 1 mM EDTA, 0.5 mM EGTA, 0.1% Na-Deoxycholate and 0.5% N-lauroylsarcosine with protease inhibitor (Lysis Buffer 3). Chromatin was then sonicated with ActiveMotif Sonicatior (amplitude: 25%, on=20s, off=30s, 20 cycles). The sonicated chromatin was centrifugated for 10 minutes, 16000rcf at 4°C. The supernatant was collected and supplemented with 10% Triton X-100. 10% of the sonicated chromatin was kept as input DNA. For each ChIP reaction, 5 μl cold lysisg antibody were added to the remaining sonicated chromatin. ChIP samples were rotated vertically at 4°C overnight (12-16 hrs) to bind antibody to chromatin. On the next day, 50-75 μl cold lysisl magnetic Dynabeads (Protein G) were washed three times (3X) in 1 mL cold Block Solution (0.5% BSA (w/v), 1x PBS). Antibody-bound chromatin was added to beads and inverted to mix. Then rotated vertically at 4°C for at least 4 hrs. Afterwards, bound beads were washed 5X in 1 mL cold RIPA buffer (50 mM Hepes, 500 mM LiCl, 1 mM EDTA, 1% NP-40, 0.7% Na-Deoxycholate). Then, samples were washed once in 1 mL TE + 50 mM NaCl on ice and centrifuged for 3 minutes at 1000rcf, 4°C to remove all remaining TE. 210μl cold lysisl Elution Buffer was added to the beads and DNA was eluted for 15 minutes at 65°C, 900 RPM. Samples were centrifuged (1 minute, 16000rcf, RT) and the supernatants were transferred (~200 uL) to fresh microfuge tubes. Both ChIP and input samples were then reverse-crosslinked and treated with RNAse and Proteinase K. Finally, DNA was extracted by phenol-chlorophorm followed by ethanol precipitation and resuspension in water. DNA content was measured using Qubit and the HS DNA Kit (Invitrogen Q32851). All antibodies used in this study have been previously reported as suitable for ChIP: H3K4me3 (39159, Active Motif), H3K27ac (39133, Active Motif), CTCF (61312, Active Motif), H3K27me3 (39155, Active Motif).

### HiChIP

HiChIPs were performed as described^33^ with some modifications. We generally replaced the ChIP protocol with the one described above, we cut and ligated overnight (NEB T4 Ligase, #M0202 instead of Invitrogen T4, 15224-041) and DNA was extracted with phenol-chlorophorm not a kit. For low cell numbers, we increased the centrifugation time to 30 minutes and 15 minutes after lysis, as well as 30 minutes after ligation to see a pellet more accurately. Generally, 12 cycles were used for Tn5 Nextera PCR amplification (Illumina Nextera DNA UD Indexes Kit). We aimed for 100M read pairs for each run on a HiSeq 2500 sequencer (Illumina), except for the AntNPC samples which were sequenced at 50M read depth. We used approx. ¼ of a 10 cm plate for each cell culture HiChIP replicate. For each murine E10.5 brain replicate, we used 1M cells.

### Generation of CRISPR/Cas9-mediated PE deletions in mice

The CRISPR/Cas9 endonuclease-mediated PE deletions were generated by the CECAD *in vivo* Research Facility (ivRF) by pronuclear injection of the Cas9 nuclease mRNA and protein, tracrRNA, and crRNAs into C57BL/6 zygotes, as previously described^84^. Cas9 nuclease (Addgene #1074181), tracrRNA (Addgene #1072532), and custom crRNA sequences were purchased from Integrated DNA Technologies (IDT; Coralville, Iowa, USA). The animals were housed and bred under standard conditions in the CECAD ivRF. The breedings described were approved by the Landesamt für Natur, Umwelt, und Verbraucherschutz Nordrhein-Westfalen (LANUV), Germany (animal application 84-02.04.2015.A405).

### Designing and molecular cloning of chicken gRNA

The template sequence flanking the PE region for each gene of interest (*Lhx5*, *Sox1* and *Sox3*) was obtained from the UCSC Genome Browser and used for gRNA design. The gRNA sequences were designed using Benchling (**Table S1**). We followed the standard principles to avoid off-target effects when choosing between multiple gRNA targets for each PE and cloned into the modified U6.3>gRNA.f+e backbone (Addgene #99139) as described in ^54^. After cloning the gRNA into the backbone, the positive bacterial clones were tested using colony PCR with the corresponding forward and reverse gRNA oligo described in the **Table S1**. The resulting PCR products were analyzed by Sanger sequencing (SeqLab). We used a control gRNA with a protospacer sequence not found in the chicken genome (GCAC-TGCTACGATCTACACC) which is already cloned into U6.3>Control.gRNA f+e vector, also provided by Prof. Dr. Marianne Bronner (Addgene #99140). To validate the CRIPSR/Cas9 targeting efficiency, PCR-based genotyping was done.

### Genotyping of PE deletions

Whole chicken embryos were electroporated with pCAGG>nls-hCas9-nls-GFP together with either U6.3>PE_*Lhx5* gRNA f+e, U6.3>PE_*Sox1* gRNA f+e or U6.3>PE_*Six3* gRNA f+e at stage HH9, then incubated until stage HH14-16 was reached. Live embryos were dissected in sterile PBS (1x) on ice and electroporated GFP-positive regions were isolated using surgical scissors. After isolation, neural tube and eye sections were pooled in a 1.5 ml tube separately for each embryo and used immediately-dissociated for genotyping with one volume of Lysis Buffer (LyB; 50 mM KCL, 10 mM TRIS pH 8.3, 2.5 mM MgCl_2,_ 0.45% NP40 and 0.45% Tween 20), containing Proteinase K (1μl cold lysisl of 20 μl cold lysisg/μl cold lysisl of Proteinase K for every 25μl cold lysisl of LyB) and incubated at 55°C for 1 hour with frequent shaking to mix. The lysate was then heated to 95°C for another 10 minutes to inactivate Proteinase K. We then tested for the presence of the PE deletions by PCR-based genotyping using the specific primers described in **Table S1**.

To detect the PE *Lhx5* deletion in mice, genomic DNA was isolated from ear punches and analyzed by PCR using the primers shown in **Table S1**. The deletion was further confirmed by Sanger sequencing of the deletion-specific PCR products (Seqlab).

### Chicken Embryos

Fertilized chicken eggs (white leghorn; Gallus gallus domesticus) were obtained from a local breeder (LSL Rhein-Main) and incubated at 37°C and 80% humidity in a normal poultry egg incubator (Typenreihe Thermo-de-Lux). Following microsurgical procedures, the eggs were re-incubated until the embryos reached the desired developmental stages. The developmental progress was determined according to the staging system of HH^85^.

### *In Ovo* Electroporation

Electroporations were performed using stage HH9 chicken embryos. 3.5–4 mL of albumin were removed by using a medical syringe to lower the blastoderm and make the embryo accessible for manipulation. The eggs were windowed, and the extra embryonic membrane was partially removed in the region to be electroporated. For knockout experiments, 5 μl cold lysisg/μl cold lysisl pCAGG>nls-hCas9-nls-GFP (Addgene #99141) and 3 μl cold lysisg/μl cold lysisl U6.3>Lhx5/Sox1/Six3 gRNA f+e was microinjected together with the Fast Green solution (Sigma) at a 2:1 ratio to ease the detection of the injection site of the developing neural tube and eye respectively with the help of borosilicate glass capillaries and electroporated as previously reported^86–88^ using the Intracel TSS20 OVODYNE Electroporator. Control embryos were electroporated with 1.5 μl cold μg/μl cold lysisl U6.3>Control.gRNA f+e along with pCAGG>nls-hCas9-nls-GFP. Following electroporation, the eggs were sealed with medical tape and re-incubated until the desired developmental stages (HH14-16) was reached.

### *In situ* hybridization (ISH)

At the desired stages, embryos were dispatched and fixed in 4% PFA/PBT overnight for ISH. Whole mount ISH of electroporated chicken embryos (HH14-HH16) and mutant/wt mice embryos (E8.5 and E9.5) was performed with probes against the target genes as previously described^88,89^. For *Lhx5*, *Sox1* and *Six3*, T7 promoter-containing PCR products were synthesized from stage HH9-HH14 chick cDNA. The gel-purified PCR products were used as templates for synthesis of antisense RNA probes using T7 polymerase enzyme. Primers are described in the **Table S1**. Riboprobes were labeled with a digoxigenin RNA labeling kit (Thermofisher, AM1324). Furthermore, mutant/wt mice embryos were tested for *Lhx5*, using DIG-labeled RNA probe against mouse *Lhx5* (986 bp) (**Table S1**) by cloning the template into the pCR™II-TOPO (Thermofischer) vector, digestion with XhoI and *in vitro* transcription using the SP6 polymerase.

### Chicken embryo sectioning and microscopy

Selected embryos were sectioned using a vibratome (Leica) at a thickness of 30-35 um. Light microscopy images were taken on a Olympus SZX16 stereomicroscope, To prepare the permanent slide, sections were embedded in Aquatex (Merck).

### ChIP-& ATAC-seq analysis

Essentially the same analysis workflows were used for ChIP-seq and ATAC-seq data. Sequencing reads were mapped with bwa-0.7.7 mem^90^, then converted from SAM to BAM (with samtools-1.2^91^) and sorted (with samtools-1.2). We then removed duplicate reads (with picard-tools-2.5.0^92^) and generated an index file (with samtools-1.2). Finally, bigWig/bedGraph files were generated with deepTools^93^ bamCoverage 2.5.7, normalized to 1x depth of coverage (reads per genome coverage), profiled and uploaded into the UCSC browser^94^ (hub hosted on cyverse^95^). Peaks are called with macs 2.1.1.20160309^96^. The narrow peak mode was used for H3K4me3, ATAC-seq/DNAse-seq and transcription factors. The broad peak mode was used for H3K27ac and H3K27me3. The Default threshold q=0.1 was used. The ChIP-seq data from mouse WT *in vitro* pluripotent cell types (i.e. serum+LIF ESC, 2i+LIF ESC and EpiLC) was obtained from Cruz-Molina et al.^*5*^ and Bleckwehl et al.^*79*^. All publically available ChIP-seq and ATAC-seq datasets used in this work, including those generated by ENCODE^97^, are listed in **Table S1**. Multiple files of the same entity were merged using bigWigMerge.

### Poised enhancer calling

#### Mouse in vitro pluripotent cell types

H3K27ac, H3K4me1 and H3K27me3 broad peaks (q**≤**0.1) in serum + LIF ESC, 2i + LIF ESC and EpiLC were additionally filtered by requiring fold-enrichments over input of at least 5, 2 and 2, respectively. The resulting peaks were extended +/− 1kb.

p300 and ATAC-seq narrow peaks (q**≤**0.1) in serum + LIF ESC, 2i + LIF ESC and EpiLC were additionally filtered by requiring fold-enrichments over input t of at least 4.

H3K27me3 and ATAC-seq/p300 peaks were intersected separately for each cell type. H3K27ac peaks of all three mouse pluripotent cell types were pooled. Poised enhancers (PEs) were called in mm10 by subtracting the H3K27ac pooled peaks from the genomic regions in which the H3K27me3 and the ATAC-seq/p300 peaks overlapped. Active enhancers were called by pooling H3K27me3 peaks from all three mouse pluripotent cell types, which were then substracted from the genomic regions in which the H3K27acAC-seq/p300 peaks overlapped. Primed enhancers were called by subtracting the pooled H3K27ac and H3K27me3 peaks from the pooled H3K4me1 peaks identified in three three mouse pluripotent cell types. For all the previous enhancer sets, genomic regions located proximal to gene TSS (+/− 5kb) were filtered out. Lastly, enhancers belonging to the same category and overlapping by at least 1bp were merged. PoiAct enhancers were either called by default by intersecting PEs with AntNPC H3K27ac peaks (q**≤** 0.1; broad; fold enrichment ≥ 5; +/− 1kb) or alternatively by intersecting distally located (>10kb from TSS) PEs with E10.5 H3K27ac ChIP-Seq^97^ peaks (q**≤** 0.1; broad; fold enrichment ≥ 1). The latter were called by intersection of called peaks in two biological replicates of fore-, mid- and hindbrain each (**Table S1**). All operations were conducted using bedtools2-2.19.0^98^.

#### Human, chicken and zebrafish

For human, chicken and zebrafish, *de novo* PEs were called using similar criteria to those described for mouse cells, but we extended each H3K27me3 and H3K27ac (only for human ESC) peak by +/− 2.5kb and we only considered regions located distally (>10kb) from gene TSS. For calling ChIP-seq and ATAC-seq peaks we always used q**≤** 0.1. Moreover for the p300/ ATAC-seq peaks we required the following fold-enrichment over input thresholds: human ≥ 3, chicken ≥ 3 and zebrafish ≥ 5. For the H3K27me3 peaks we required the following fold-enrichment over input thresholds: human ≥ 3, chicken ≥ 1 and zebrafish ≥ 3. The H3K27ac peaks in human were required to have fold-enrichment thresholds ≥3.

All generated enhancer bed files are listed in **Table S2**.

PE sets were annotated using GREAT 3.0^79^, except for the chicken PE, which were annotated with ConsensusPathDb^80^. Zebrafish *de novo* PE had to be lifted to assembly danRer7 for annotation with GREAT. The used assemblies and annotations are listed in **Table S1**.

### Conservation analysis

All available UCSC vertebrates were considered if they had an available liftOver chain file from mm10 (**Table S1** for all used genome builds). Specific genome build versions were used to match Ensembl TSS annotations of the species in which Bio-CAP data was available^22^. Identity thresholds of 50% were used.

All species depictions are royalty-free / limited licensed under the stock IDs 57341071, 1666821223, 750090523, 99158564, 182837423, 194504681 & 1536755681.

### Non-methylated islands (NMI)

NMI and Bio-CAP data was obtained from Long et al.^22^. Processed NMI bed files were initially used to calculate NMI to PE distances. Raw reads were downloaded from GEO^99^ (GSE43512), aligned using bowtie2-2.2.0^100^ and profiled using deepTools bamCoverage 2.5.7. Narrow NMI peaks were called using macs 2.1.1.20160309 *callpeak (*q=0.01; “--extsize 300 --mfold 10 30”).

### HiChIP

Except for murine E10.5 brain samples, biological/technical replicates were merged. Reads were aligned and quality-controlled through HiC-Pro 2.10^101^. Initial anchor calling was conducted using macs 2.1.1.20160309 *callpeak* on bowtie alignments with default q=0.1. The resulting peaks were passed to FitHiChIP 7.0^81^ (29th April 2019) for final loop calling (in L_COV mode). Basic loop quality control statistics including distance plots were conducted using diffloop^102^. hichipper 0.7^103^ was used to generate bedgraph files and ChIP signal profiles. BigWig files were generated after clipping bedGraph files using bedClip. Final tracks were visualized in WashU^104^ or UCSC genome browsers. Loop anchors were overlapped with TSS and annotated using ConsensusPathDb^80^. Hi-C matrices were processed and normalized using cooler^105^ (5kb resolution), then plotted with coolpup.py^106^ using default settings with “--padding 100”. HiChIP samples were coverage-normalized (unbalanced) and Hi-C samples were KR-balanced. Loopiness values were calculated on the single center pixel normalized to corners (“--enrichment 1 --norm_corners 1”). Single loci were visualized using HiCPlotter^107^ with standard RefSeq annotations. PcG domains for looping analyses were called by overlapping significant peaks (macs2; narrow; q=0.1) of EED^108^ and RING1b^109^ ChIP-seq binding sites. TAD evaluations were conducted using coordinates^110^ included within the HiCPlotter package lifted over from mm9 to mm10. All residual statistical analyses were conducted using custom-made scripts in R (3.6.0; 2019-04-26).

### RNA-seq and scRNA-seq

The expression of targets genes (evaluated through sign. E14 H3K27me3 HiChIP interactions) for different poised and PoiAct enhancer sets was plotted for AntNPC and mESC RNA-seq^5^, as well as for blastulation, gastrulation and neurulation samples from two scRNA-Seq data sets^111,112^. Expression differences between differentiation or developmental stages were evaluated using Wilcoxon tests. Processed data was obtained from the respective GEO entries. To circumvent batch effects in both plots, expression was divided by housekeeping genes (all available eukaryotic translation elongation factors and actin molecules) mean expression.

### Data and Software Availability

All ChIP-seq, ATAC-seq and HiChIP data sets generated in this work have been deposited into the GEO repository under accession number GEO: GSE160657.

All public data sets used are listed in **Table S1**.

---

**Fig. S1:**
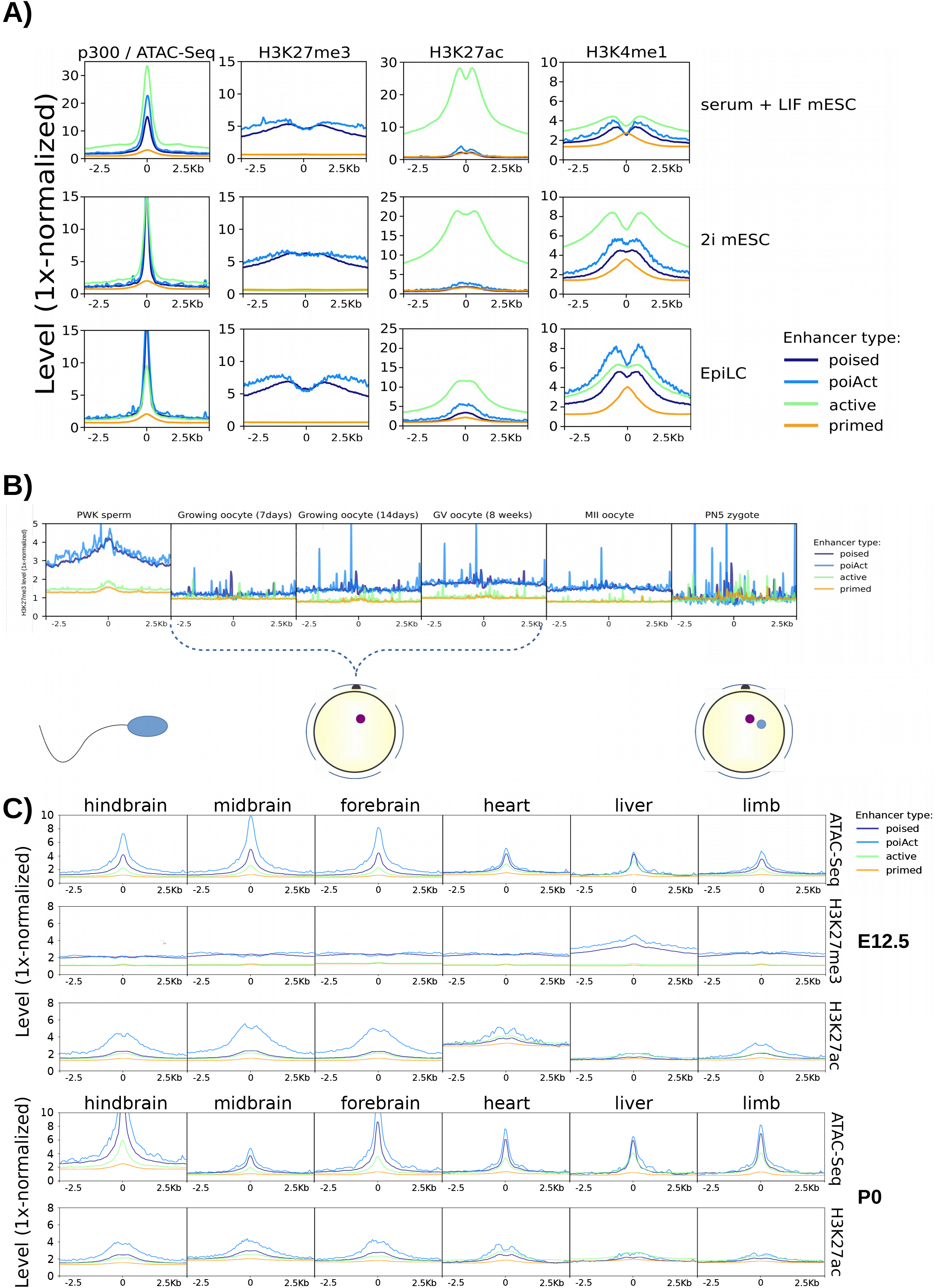
Epigenomic features of PEs during mouse embryogenesis. **(A)** H3K27ac, H3K4me1 and H3K27me3 signals from serum+LIF mESC, 2i mESC and EpiLC^1^ are shown around poised, active, primed and PoiAct enhancers identified in *in vitro* pluripotent cells (**Fig. 1; Methods**). ATAC-seq signals are shown for 2i mESC and EpiLC and p300 signals for serum+LIF mESC. **(B)** H3K27me3 ChIP-Seq profiles generated in male and female mouse germ cells^2^ are shown around poised, active, primed and PoiAct enhancers identified in *in vitro* pluripotent cells. **(C)** ATAC-seq, H3K27me3 and H3K27ac signals from six different mouse embryonic (E12.5; upper panel) or postnatal (P0; lower panels) tissues are shown around poised, active, primed and PoiAct enhancers identified in *in vitro* pluripotent cells.

**Fig. S2:**
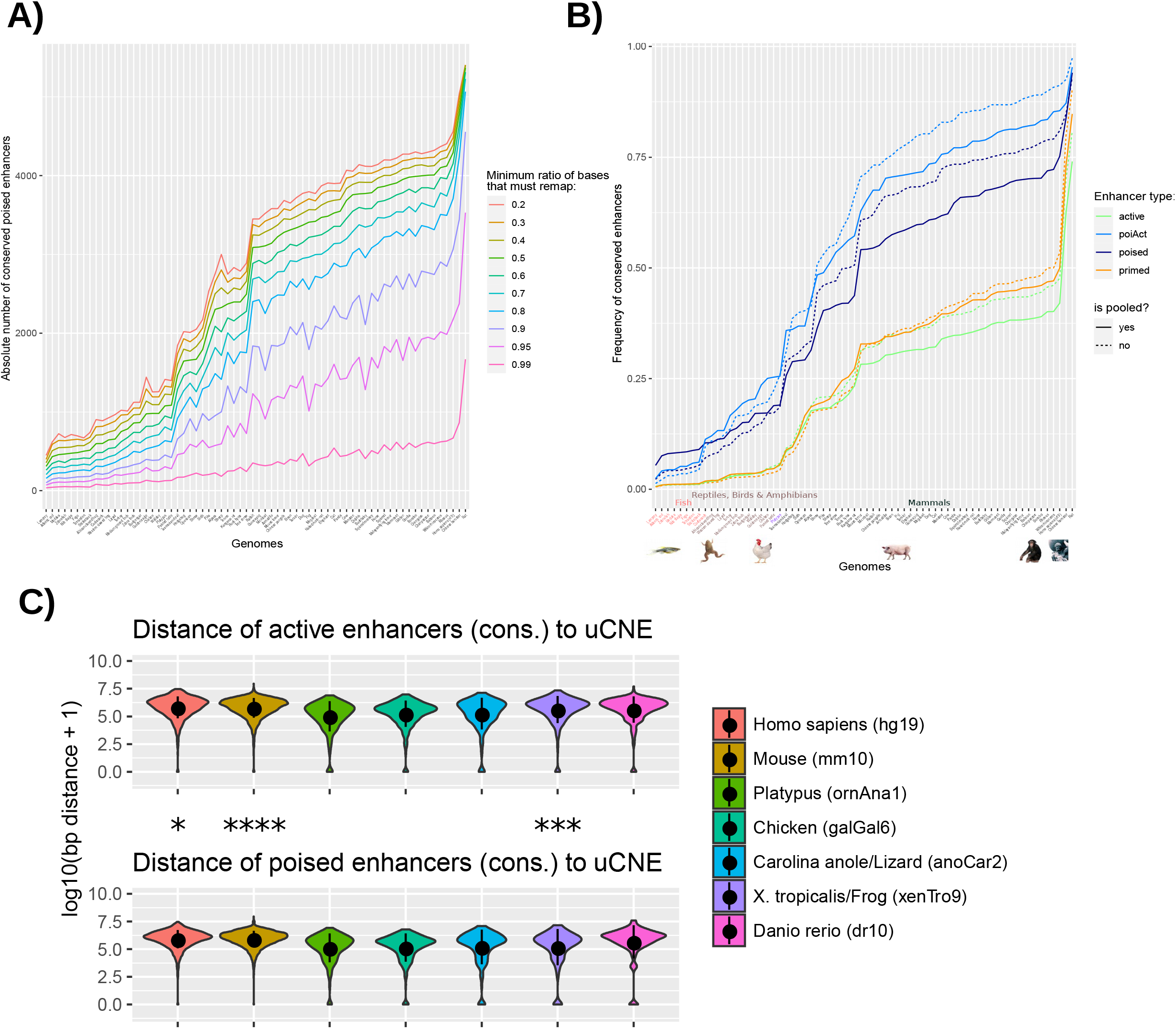
Conservation analysis of mouse PE across vertebrates. **(A)** Absolute number of mouse PEs conserved across 68 vertebrates (**Table S1**) using different mappability thresholds (mapping ratios 0.2-0.99; color legend). **(B)** Sequence conservation across 68 vertebrate species was measured for poised, active, primed and PoiAct enhancers identified in mouse pluripotent cells using a mappability threhold of 0.5. The previous enhancer sets were further divided depending on whether they were called considering several pluripotent cell types as defined in **Fig. 1** („pooled“; solid lines) or only in serum+LIF mESC as originally described^*3*^ (dashed lines). **(C)** Distance between active (upper panel) or poised (lower panel) mouse enhancers conserved in the indicated vertebrate species and ultraconserved non-coding elements (uCNE) identified in the same species^4^. The asterisks indicate that the distances between either poised or active enhancers and uCNU are significantly different (Wilcoxon test). Human and frog PE (FC=1.15, p=0.012; FC=1.39, p=0.00036; Wilcoxon test) are closer to uCNE, but mouse and zebrafish PE (FC=0.89, p=1.24e-16; FC=0.62, p=0.075; Wilcoxon test) are more distal to uCNE when compared to active conserved enhancers. * p ≤ 0.05, ** p ≤ 0.01, *** p ≤ 0.001, **** p ≤ 0.0001.

**Fig. S3:**
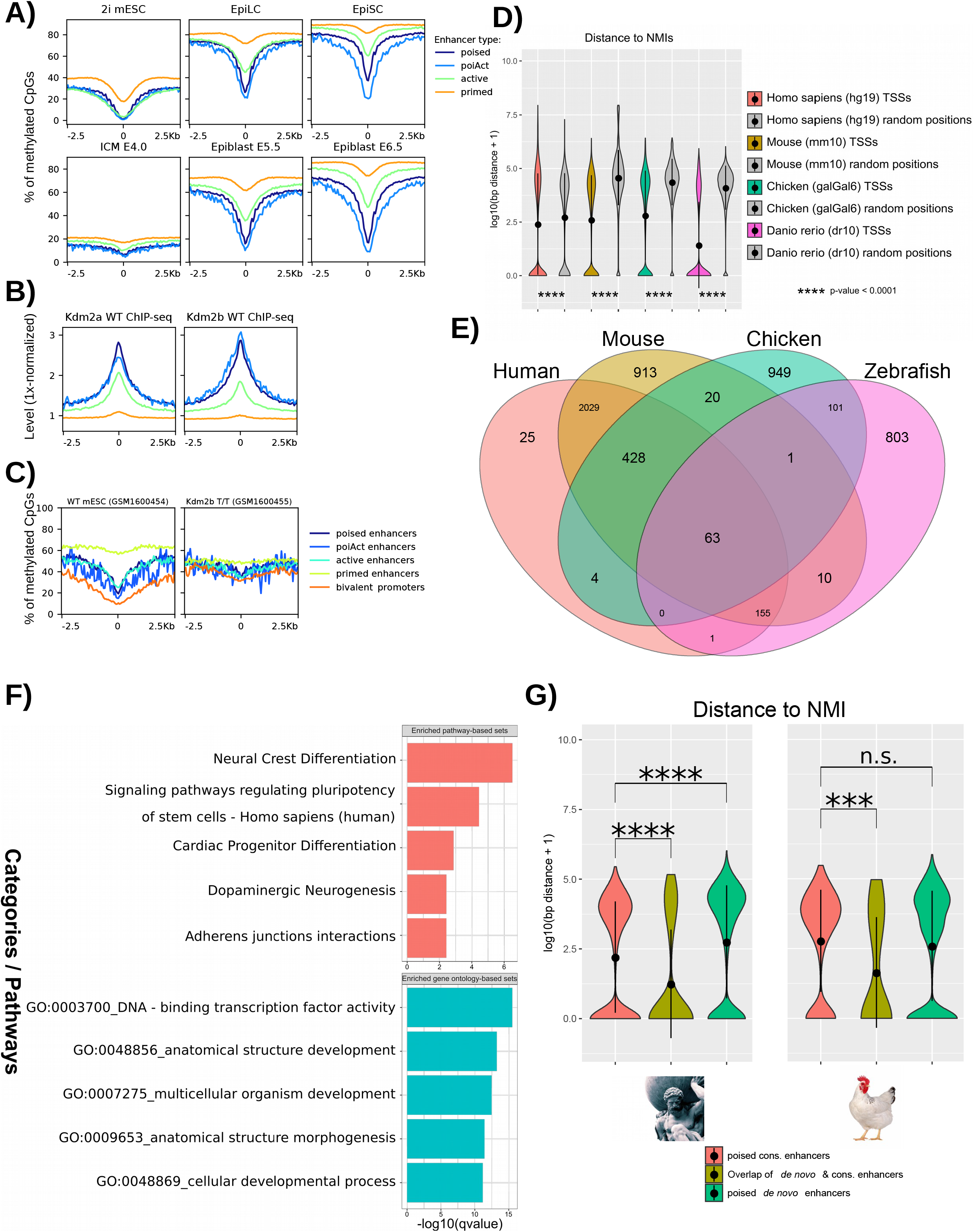
Genetic, epigenetic and regulatory features of PEs are conserved across vertebrates. **(A)** Whole-genome bisulfite sequencing (WGBS) / STEM-seq data generated in either *in vitro* (2i ESC, EpiLC and EpiSC; upper row) or *in vivo* (E4.0 ICM, E5.5 epiblast and E6.5 epiblast; lower row) mouse pluripotent cells^5–7^ was analysed in order to show CpG methylation levels around poised, active, primed and PoiAct enhancers identified in *in vitro* pluripotent cells (**Fig. 1; Methods**). **(B)** ChIP-seq signals for KDM2A and KDM2B in mESC are shown around poised, active, primed and PoiAct enhancers identified in *in vitro* pluripotent cells. **(C)** CpG methylation levels as measured by Reduced-Representation Bisulfite Sequencing (RRBS) in WT mESC and *Kdm2b*^−/−^ mESC^8^ are shown around bivalent promoters and poised, active, primed and PoiAct enhancers identified in *in vitro* pluripotent cells. CpG methylation levels increased significantly in *Kdm2b*^−/−^ mESC compared to WT mESC at PE (FC=2.43; p < 2.2e-16; Wilcoxon test), bivalent promoters (FC=3.59; p < 2.2e-16; Wilcoxon test) and, to a lesser degree, at active enhancers (FC=1.41; p < 2.2e-16; Wilcoxon test). **(D)** The quality of the NMIs identified by Bio-CAP in different vertebrates is illustrated by the significantly shorter distances separating the NMIs from TSS than from 1000 randomly-sampled genomic regions in each of the investigated species. **(E)** All the *de novo* PEs identified in different vertebrate species (including the mouse PEs described in **Fig. 1** and located >10kb from TSS) were linked to the nearest gene. Then, homologous genes were retrieved from biomaRt^9^ and overlaps between species were calculated with Venn diagrams. **(F)** Pathway and Gene Ontology analysis for the overlapping genes (n=63) shown in (E) were calculated with ConsensusPathDB. **(G)** Distances to NMIs are shown for different PE categories identified in either humans (left panel) or chicken (right panel). The *de novo* PE overlapping with conserved mouse PE (human: n=183, chicken: n=49) were the most proximal to NMIs. * p ≤ 0.05, ** p ≤ 0.01, *** p ≤ 0.001, **** p ≤ 0.0001.

**Fig. S4:**
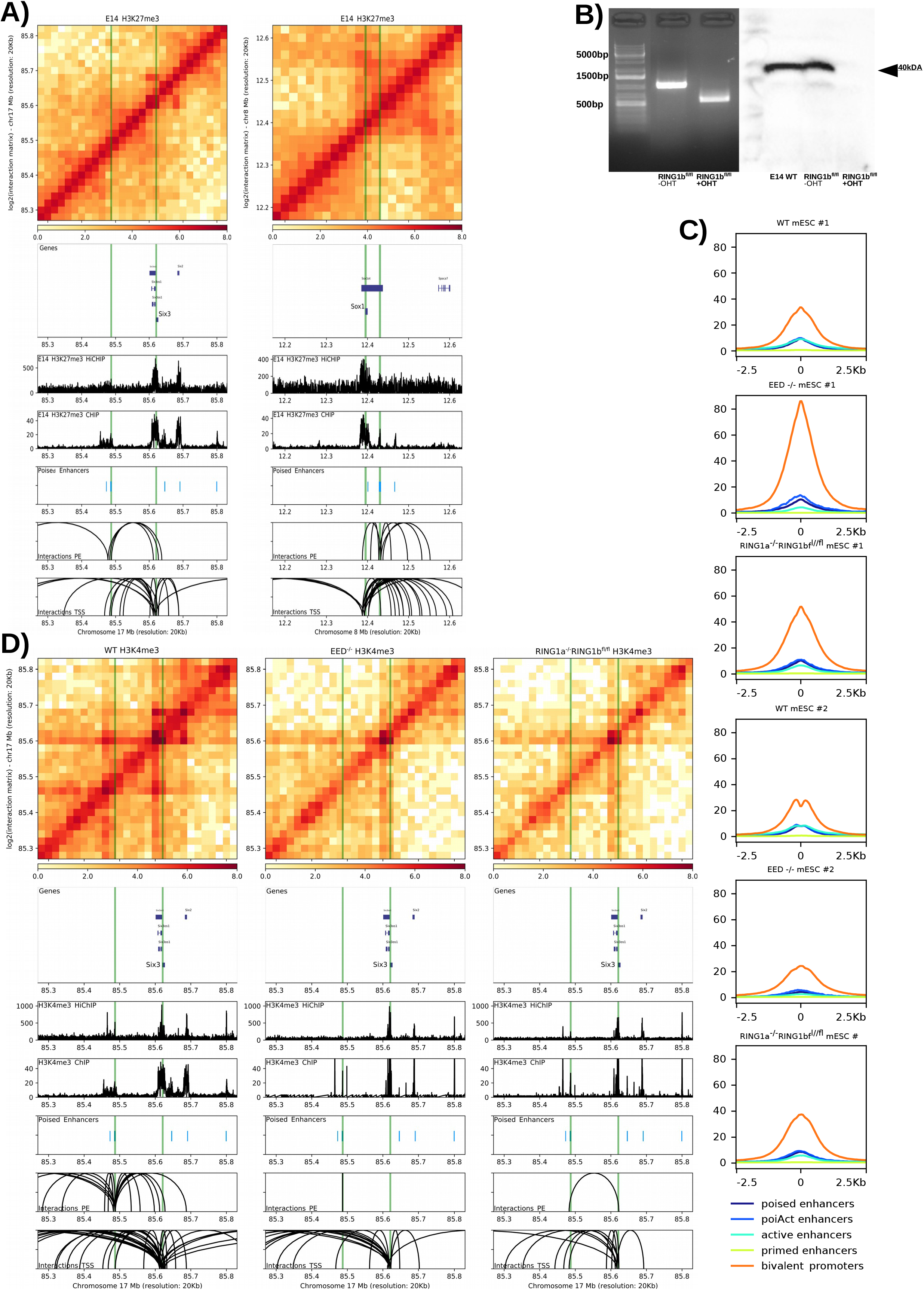
The interactions between PEs and bivalent promoters are globally mediated by PRC1. **(A)** H3K27me3 HiChIP and ChIP-seq profiles generated in mESC are shown around the *Six3* (left) and *Sox1* (right) loci. Both the *Six3 and Sox1* TSS as well as the PEs previously shown to control these genes in AntNPC (PE *Six3* (−133kb) and PE *Sox1* (+35kb)) are marked with green vertical lines. <u>Upper panels:</u> heatmap of log2 interaction intensities based on H3K27me3 HiChIP data generated in mESC. <u>Two medium panels:</u> ChIP-seq and 1D HiChIP signals for H3K27me3 in mESC. <u>Lower two panels:</u> significant (p<0.01; FitHiChIP) interactions were called using the H3K27me3 HiChIP PETs (paired-end tags) and loops in which one of the anchors overlaps (+/− 10kb) either the PE or the TSS are shown. **(B)** *Ring1a*^−/−^*Ring1b*^*fl*/*fl*^ mESC were either left untreated or treated with tamoxifen for 72 hours. The loss of Ring1b after tamoxifen treatment was confirmed by PCR genotyping (left) and Western blot analysis (right). The PCR primers used for detecting the *Ring1b* deletion are shown in **Table S1**. **(C)** ChIP-seq data for H3K4me3 was generated as biological replicates in WT mESC, *Eed*^−/−^ mESC and Tamoxifen-treated *Ring1a*^−/−^*Ring1b*^*fl*/*fl*^ mESC. H3K4me3 profiles are shown around bivalent promoters and poised, active, primed and PoiAct enhancers identified in *in vitro* mouse pluripotent cells. **(D)** H3K4me3 HiChIP in WT, *Eed*^−/−^ mESC and Tamoxifen-treated *Ring1a*^−/−^*Ring1b*^*fl*/*fl*^ mESC around the *Six3* locus.

**Fig. S5:**
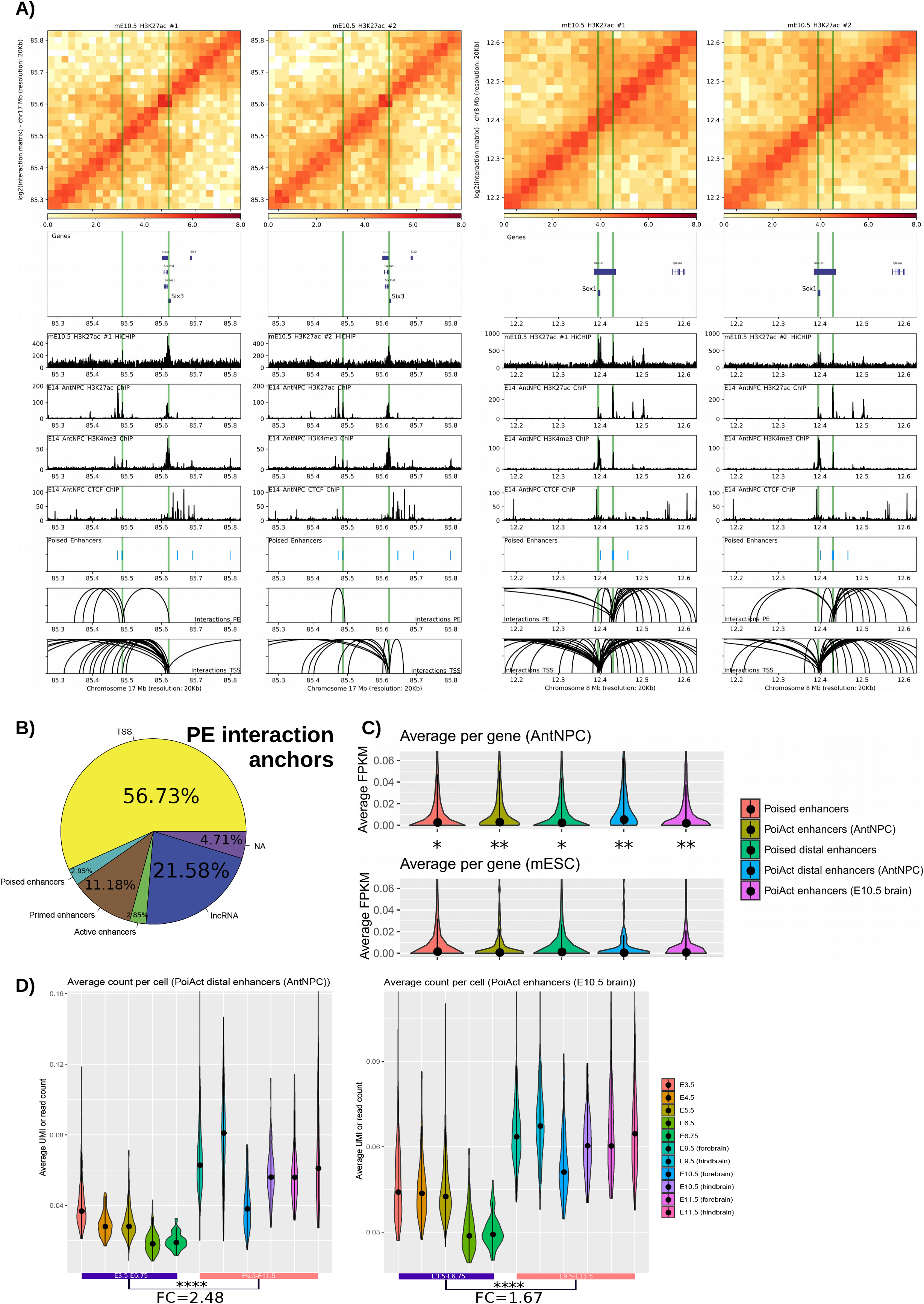
Physical interactions between PEs and their target genes in the embryonic mouse brain. **(A)** H3K27ac HiChIP and ChIP-seq profiles generated in E10.5 mouse brains and AntNPC, respectively, are shown around the *Six3* (left) and *Sox1* (right) loci. Both the *Six3 and Sox1* TSS as well as the PEs previously shown to control these genes in AntNPC (PE *Six3* (−133kb) and PE *Sox1* (+35kb)) are marked with green vertical lines. <u>Upper panels:</u> heatmap of log2 interaction intensities based on H3K27ac HiChIP data generated in E10.5 mouse brain. <u>Two medium panels:</u> 1D HiChIP and ChIP-seq signals for H3K27ac, H3K4me3 and CTCF in E10.5 mouse brain and AntNPC, respectively. <u>Lower two panels:</u> significant (p<0.01; FitHiChIP) interactions were called using the H3K27ac HiChIP PETs (paired-end tags) from E10.5 mouse brain and loops in which one of the anchors overlaps (+/−10kb) either the PEs or the TSS are shown. **(B)** Significant (p<0.01; FitHiChIP) interactions were called using the H3K27ac HiChIP PETs from E10.5 mouse brain. Interaction anchors were overlapped with PEs and their interaction partners were hierarchically annotated with the categories shown in the pie chart (**Methods**). **(C)** The target genes of different mouse PE subsets were defined using the significant H3K27me3 HiChIP interactions called in mESC (**Fig. 3**). Then the expression of the genes associated with each PE category was plotted using RNA-seq data generated in AntNPC and mESC. Each data point is the expression (in FPKM) of a target gene. The asterisks indicate significantly higher expression in AntNPC than in ESC (Wilcoxon test). The *PoiAct enhancers (E10.5 brain)* category was defined by overlapping mouse PEs with H3K27ac ChIP-Seq peaks (broad; q<0.1) called in E10.5 mouse fore-, mid- and hindbrain (**Table S1**). Value range was zoomed in to lower and upper IQR (interquartile range) to circumvent excessive outliers. **(D)** The expression of the genes associated with the *PoiAct distal enhancers (AntNPC)* (left) and *PoiAct enhancers (E10.5 brain)* (right) groups described in (C) was plotted using scRNA-Seq data generated during mouse embryogenesis. Each data point is the mean expression (in RPM (reads per million) for E3.5-E6.75 and TPM (transcripts per million) for E9.5-E11.5) of all target genes in one cell. The asterisks indicate significantly higher expression in E9.5-E11.5 brain than in E3.5-E6.75 embryos. Value range was zoomed in to lower and upper IQR to circumvent excessive outliers. * p ≤ 0.05, ** p ≤ 0.01, *** p ≤ 0.001, **** p ≤ 0.0001.

**Fig. S6:**
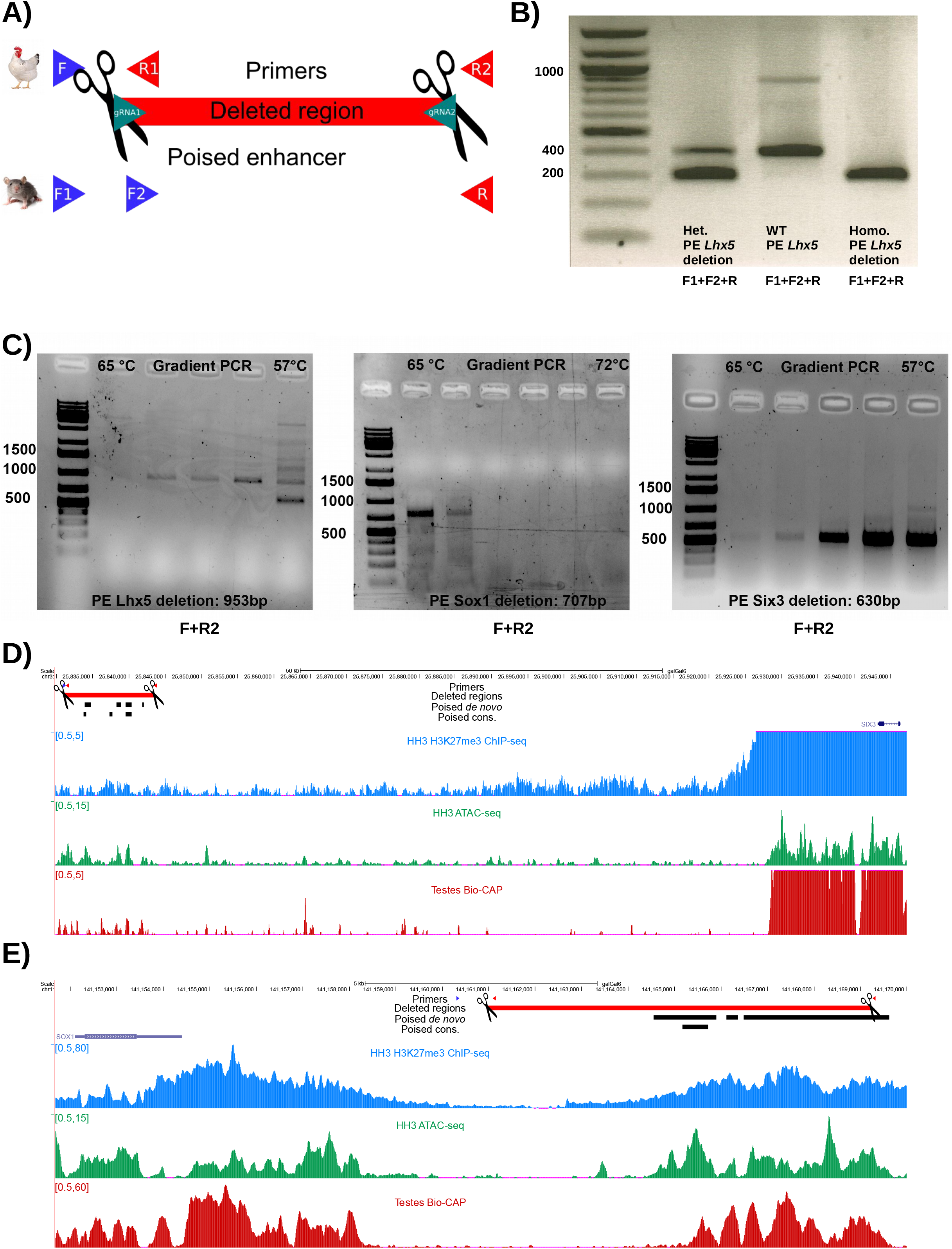
PE deletions in mouse and chicken embryos. **(A)** Scheme for genotyping of chicken embryos and transgenic mice obtained through CRISPR-Cas9 targeting of PEs. We used one or two forward primers (blue) and one or two reverse primers (red; **Table S1**) specific for the deleted and unperturbed region respectively. Primer positions for chicken are further depicted as tracks in **D**, **E** and **Fig. 6E**. **(B)** Genotyping results for the *PE Lhx5(−109kb)* deletion in transgenic mice. Used primer combinations are indicated below the gel. **(C)** The CRISPR/Cas9-targeted PEs for *Lhx5*, *Six3* and *Sox1* were PCR-amplified from electroporated chicken genomic DNA using primers specific for the deleted regions. Used primer combinations are indicated below the gel. **(D)** ATAC-seq and H3K27me3 ChIP-seq profiles generated in HH3 chicken epiblast are shown around the *Six3* locus. In addition, Bio-CAP profiles generated in chicken testes are also shown to illustrate the location of CGI/NMI in the chicken genome. The genomic region deleted in chicken embryos that includes the mouse PE *Six3* (−133kb) conserved in chicken is highlighted in red. **(E)** Similar profiles as in (E), but for the mouse PE *Sox1* (+35kb) conserved in chicken.

## References

1. Rada-Iglesias, A. et al. A unique chromatin signature uncovers early developmental enhancers in humans. Nature. 2011 Feb 10;470(7333):279–83.

2. Creyghton, M.P. et al. Histone H3K27ac separates active from poised enhancers and predicts developmental state. Proc Natl Acad Sci U S A. 2010 Dec 14;107(50):21931–6.

3. Entrevan, M., Schuettengruber, B., & Cavalli, G. Regulation of Genome Architecture and Function by Polycomb Proteins. Trends Cell Biol. 2016 Jul;26(7):511–525.

4. Schoenfelder, S. et al. Polycomb repressive complex PRC1 spatially constrains the mouse embryonic stem cell genome. Nat Genet. 2015 Oct;47(10):1179–1186.

5. Cruz-Molina, S. et al. PRC2 Facilitates the Regulatory Topology Required for Poised Enhancer Function during Pluripotent Stem Cell Differentiation. Cell Stem Cell. 2017 May 4;20(5):689–705.e9.

6. Illingworth, R.S. et al. Orphan CpG islands identify numerous conserved promoters in the mammalian genome. PLoS Genet. 2010 Sep 23;6(9):e1001134

7. van Heeringen, S.J. et al. Principles of nucleation of H3K27 methylation during embryonic development. Genome Res. 2014 Mar;24(3):401–10.

8. Li, Y. et al. Genome-wide analyses reveal a role of Polycomb in promoting hypomethylation of DNA methylation valleys. Genome Biol. 2018 Feb 8;19(1):18.

9. Pachano, T. et al. Orphan CpG islands boost the regulatory activity of poised enhancers and dictate the responsiveness of their target genes. bioRxiv. 2020.

10. Hackett, J.A. & Surani, M.A. Regulatory principles of pluripotency: from the ground state up. Cell Stem Cell. 2014 Oct 2;15(4):416–430.

11. Morgani, S., Nichols, J. & Hadjantonakis, A. The many faces of Pluripotency: in vitro adaptations of a continuum of in vivo states. BMC Dev Biol. 2017 Jun 13;17(1):7.

12. Wu, J. et al. The landscape of accessible chromatin in mammalian preimplantation embryos. Nature. 2016 Jun 30;534(7609):652–7.

13. Liu, X. et al. Distinct features of H3K4me3 and H3K27me3 chromatin domains in pre-implantation embryos. Nature. 2016 Sep 22;537(7621):558–562.

14. Zheng, H. et al. Resetting Epigenetic Memory by Reprogramming of Histone Modifications in Mammals. Mol Cell. 2016 Sep 15;63(6):1066–79.

15. Maezawa, S. et al. Polycomb protein SCML2 facilitates H3K27me3 to establish bivalent domains in the male germline. Proc Natl Acad Sci U S A. 2018 May 8;115(19):4957–4962.

16. Hiller, M. et al. Computational methods to detect conserved non-genic elements in phylogenetically isolated genomes: application to zebrafish. Nucleic Acids Res. 2013 Aug;41(15):e151.

17. Bejerano, G. et al. Ultraconserved elements in the human genome. Science. 2004 May 28;304(5675):1321–5.

18. Dickel, D.E. et al. Ultraconserved Enhancers Are Required for Normal Development. Cell. 2018 Jan 25;172(3):491–499.e15.

19. Elnitski, L. & Ovcharenko, I. The hypothesis of ultraconserved enhancer dispensability overturned. Genome Biol. 2018 May 8;19(1):57.

20. Romero, I.G. et al. A panel of induced pluripotent stem cells from chimpanzees: a resource for comparative functional genomics. Elife 2015 Jun 23;4:e07103.

21. Kaaij, L.J.T. et al. Enhancers reside in a unique epigenetic environment during early zebrafish development. Genome Biol. 2016 Jul 5;17(1):146.

22. Long, H.K. et al. Epigenetic conservation at gene regulatory elements revealed by non-methylated DNA profiling in seven vertebrates. Elife. 2013 Feb 26;2:e00348.

23. He, J. et al. Kdm2b maintains murine embryonic stem cell status by recruiting PRC1 complex to CpG islands of developmental genes. Nat Cell Biol. 2013 Apr;15(4):373–84.

24. Blackledge, N.P. et al. Variant PRC1 complex-dependent H2A ubiquitylation drives PRC2 recruitment and polycomb domain formation. Cell. 2014 Jun 5;157(6):1445–1459.

25. Chen, S., Jiao, L., Liu, X.,Yang, X. & Liu, X. A Dimeric Structural Scaffold for PRC2-PCL Targeting to CpG Island Chromatin. Mol Cell. 2020 Mar 19;77(6):1265–1278.e7.

26. Habibi, E. et al. Whole-genome bisulfite sequencing of two distinct interconvertible DNA methylomes of mouse embryonic stem cells. Cell Stem Cell. 2013 Sep 5;13(3):360–9.

27. Zylicz, J.J. et al. Chromatin dynamics and the role of G9a in gene regulation and enhancer silencing during early mouse development. Elife. 2015 Nov 9;4:e09571.

28. Farcas, A.M. et al. KDM2B links the Polycomb Repressive Complex 1 (PRC1) to recognition of CpG islands. Elife 2012 Dec 18;1:e00205.

29. Boulard, M., Edwards, J.R. & Bestor, T.H. FBXL10 protects Polycomb-bound genes from hypermethylation. Nat Genet. 2015 May;47(5):479–85.

30. Rada-Iglesias, A. et al. Epigenomic annotation of enhancers predicts transcriptional regulators of human neural crest. Cell Stem Cell. 2012 Nov 2;11(5):633–48.

31. Belaghzal, H., Dekker, J. & Gibcus, J.H. Hi-C 2.0: An optimized Hi-C procedure for high-resolution genome-wide mapping of chromosome conformation. Methods. 2017 Jul 1;123:56–65.

32. Bonev, B. et al. Multiscale 3D Genome Rewiring during Mouse Neural Development. Cell. 2017 Oct 19;171(3):557–572.e24.

33. Mumbach, M.R. et al. HiChIP: efficient and sensitive analysis of protein-directed genome architecture. Nat Methods. 2016 Nov;13(11):919–922.

34. Kundu, S. et al. Polycomb Repressive Complex 1 Generates Discrete Compacted Domains that Change during Differentiation. Mol Cell. 2017 Feb 2;65(3):432–446.e5.

35. McLaughlin, K. et al. DNA Methylation Directs Polycomb-Dependent 3D Genome Re-organization in Naive Pluripotency. Cell Rep. 2019 Nov 12;29(7):1974–1985.e6.

36. Zhang, Y. et al. Dynamic epigenomic landscapes during early lineage specification in mouse embryos. Nat Genet. 2018 Jan;50(1):96–105.

37. Joshi, O. et al. Dynamic Reorganization of Extremely Long-Range Promoter-Promoter Interactions between Two States of Pluripotency. Cell Stem Cell. 2015 Dec 3;17(6):748–757.

38. Gentile, C. et al. PRC2-Associated Chromatin Contacts in the Developing Limb Reveal a Possible Mechanism for the Atypical Role of PRC2 in HoxA Gene Expression. Dev Cell. 2019 Jul 22;50(2):184–196.e4.

39. Loubiere, V. et al. Widespread activation of developmental gene expression characterized by PRC1-dependent chromatin looping. Sci Adv. 2020 Jan 10;6(2):eaax4001.

40. Schoeftner, S. et al. Recruitment of PRC1 function at the initiation of X inactivation independent of PRC2 and silencing. EMBO J. 2006 Jul 12;25(13):3110–22.

41. Endoh, M. et al. Polycomb group proteins Ring1A/B are functionally linked to the core transcriptional regulatory circuitry to maintain ES cell identity. Development. 2008 Apr;135(8):1513–24.

42. Leeb, M. et al. Polycomb complexes act redundantly to repress genomic repeats and genes. Genes Dev. 2010 Feb 1;24(3):265–76.

43. Ferrari, K.J. et al. Polycomb-dependent H3K27me1 and H3K27me2 regulate active transcription and enhancer fidelity. Mol Cell. 2014 Jan 9;53(1):49–62.

44. Isono, K. et al. SAM domain polymerization links subnuclear clustering of PRC1 to gene silencing. Dev Cell. 2013 Sep 30;26(6):565–77.

45. Plys, A.J. et al. Phase separation of Polycomb-repressive complex 1 is governed by a charged disordered region of CBX2. Genes Dev. 2019 Jul 1;33(13-14):799–813.

46. Tatavosian, R. et al. Nuclear condensates of the Polycomb protein chromobox 2 (CBX2) assemble through phase separation. J Biol Chem. 2019 Feb 1;294(5):1451–1463.

47. Pachano, T., Crispatzu, G. & Rada-Iglesias, A. Polycomb proteins as organizers of 3D genome architecture in embryonic stem cells. Brief Funct Genomics. 2019 Nov 19;18(6):358–366.

48. Kent, S. et al. Phase-Separated Transcriptional Condensates Accelerate Target-Search Process Revealed by Live-Cell Single-Molecule Imaging. Cell Rep. 2020 Oct 13;33(2):108248.

49. Schuettengruber, B., Bourbon, H., Di Croce, L. & Cavalli, G. Genome Regulation by Polycomb and Trithorax: 70 Years and Counting. Cell. 2017 Sep 21;171(1):34–57

50. Xiang, Y. et al. Epigenomic analysis of gastrulation identifies a unique chromatin state for primed pluripotency. Nat Genet. 2020 Jan;52(1):95–105.

51. Rhodes, J.D.P. et al. Cohesin Disrupts Polycomb-Dependent Chromosome Interactions in Embryonic Stem Cells. Cell Rep. 2020 Jan 21;30(3):820–835.e10

52. Nora, E.P. et al. Targeted Degradation of CTCF Decouples Local Insulation of Chromosome Domains from Genomic Compartmentalization. Cell. 2017 May 18;169(5):930–944.e22.

53. Kishi, Y. & Gotoh, Y. Regulation of Chromatin Structure During Neural Development. Front Neurosci. 2018 Dec 11;12:874.

54. Gandhi, S., Piacentino, M.L., Vieceli, F.M. & Bronner, M.E. Optimization of CRISPR/Cas9 genome editing for loss-of-function in the early chick embryo. Dev Biol. 2017 Dec 1;432(1):86–97.

55. Carl, M., Loosli, F. & Wittbrodt, F. Six3 inactivation reveals its essential role for the formation and patterning of the vertebrate eye. Development. 2002 Sep;129(17):4057–63.

56. Kikuta, H. et al. Genomic regulatory blocks encompass multiple neighboring genes and maintain conserved synteny in vertebrates. Genome Res. 2007 May;17(5):545–55.

57. Akalin, A. et al. Transcriptional features of genomic regulatory blocks. Genome Biol. 2009;10(4):R38.

58. Harmston, N. et al. Topologically associating domains are ancient features that coincide with Metazoan clusters of extreme noncoding conservation. Nat Commun. 2017 Sep 5;8(1):441.

59. Mahmoudi, T., Katsani, K.R. & Verrijzer, C.P. GAGA can mediate enhancer function in trans by linking two separate DNA molecules. EMBO J. 2002 Apr 2;21(7):1775–81.

60. Calhoun, V.C., Stathopoulos, A. & Levine, M. Promoter-proximal tethering elements regulate enhancer-promoter specificity in the Drosophila Antennapedia complex. Proc Natl Acad Sci U S A. 2002 Jul 9;99(14):9243–7.

61. Calhoun, V.C. & Levine, M. Long-range enhancer-promoter interactions in the Scr-Antp interval of the Drosophila Antennapedia complex. Proc Natl Acad Sci U S A. 2003 Aug 19;100(17):9878–83.

62. Engström, P.G., Ho Sui, S.J., Drivenes, O., Becker, T.S. & Lenhard, B. Genomic regulatory blocks underlie extensive microsynteny conservation in insects. Genome Res. 2007 Dec;17(12):1898–908.

63. Perino, M. et al. MTF2 recruits Polycomb Repressive Complex 2 by helical-shape-selective DNA binding. Nat Genet. 2018 Jul;50(7):1002–1010.

64. Boyle, S. et al. A central role for canonical PRC1 in shaping the 3D nuclear landscape. Genes Dev. 2020 Jul 1;34(13-14):931–949.

65. Cao, R. et al. Role of histone H3 lysine 27 methylation in Polycomb-group silencing. Science. 2002 Nov 1;298(5595):1039–43.

66. Blackledge, N.P. & Klose, R. CpG island chromatin: a platform for gene regulation. Epigenetics. 2011 Feb;6(2):147–52.

67. Deaton, A.M. & Bird, A. CpG islands and the regulation of transcription. Genes Dev. 2011 May 15;25(10):1010–22.

68. Wachter, E. et al. Synthetic CpG islands reveal DNA sequence determinants of chromatin structure. Elife. 2014 Sep 26;3:e03397.

69. Mas, G. et al. Promoter bivalency favors an open chromatin architecture in embryonic stem cells. Nat Genet. 2018 Oct;50(10):1452–1462.

70. Kondo, T. et al. Polycomb potentiates meis2 activation in midbrain by mediating interaction of the promoter with a tissue-specific enhancer. Dev Cell. 2014 Jan 13;28(1):94–101.

71. Pu, L. & Sung, Z.R. PcG and trxG in plants - friends or foes. Trends Genet. 2015 May;31(5):252–62.

72. Fudenberg, G. et al. Formation of Chromosomal Domains by Loop Extrusion. Cell Rep. 2016 May 31;15(9):2038–49.

73. Schwarzer, W. et al. Two independent modes of chromatin organization revealed by cohesin removal. Nature. 2017 Nov 2;551(7678):51–56.

74. Rao, S.S.P. et al. Cohesin Loss Eliminates All Loop Domains. Cell. 2017 Oct 5;171(2):305–320.e24.

75. Perry, M.W., Boettiger, A.N., Bothma, J.P. & Levine, M. Shadow enhancers foster robustness of Drosophila gastrulation. Curr Biol. 2010 Sep 14;20(17):1562–7.

76. Cannavò, E. et al. Shadow Enhancers Are Pervasive Features of Developmental Regulatory Networks. Curr Biol. 2016 Jan 11;26(1):38–51.

77. Osterwalder, M. et al. Enhancer redundancy provides phenotypic robustness in mammalian development. Nature. 2018 Feb 8;554(7691):239–243.

78. Minoux, M. et al. Gene bivalency at Polycomb domains regulates cranial neural crest positional identity. Science. 2017 Mar 31;355(6332):eaal2913.

79. Bleckwehl, T. et al. Enhancer priming by H3K4 methylation safeguards germline competence. bioRxiv. 2020.

80. McLean, C.Y. et al. GREAT improves functional interpretation of cis-regulatory regions. Nat. Biotechnol. 2010 28(5):495–501.

81. Herwig, R., Hardt, C., Lienhard, M. & Kamburov, A. Analyzing and interpreting genome data at the network level with ConsensusPathDB. Nat Protoc. 2016 Oct;11(10):1889–907.

82. Bhattacharyya, S., Chandra, V., Vijayanand, P. & Ay, F. Identification of significant chromatin contacts from HiChIP data by FitHiChIP. Nat Commun. 2019 Sep 17;10(1):4221.

83. Buenrostro, J.D., Giresi, P.G., Zaba, L.C., Chang, H.Y. & Greenleaf, W.J. Transposition of native chromatin for fast and sensitive epigenomic profiling of open chromatin, DNA-binding proteins and nucleosome position. Nat Methods. 2013 Dec;10(12):1213–8.

84. Chu, V.T. et al. Efficient CRISPR-mediated mutagenesis in primary immune cells using CrispRGold and a C57BL/6 Cas9 transgenic mouse line. Proc Natl Acad Sci U S A. 2016 Nov 1;113(44):12514–12519.

85. Hamburger, V. & Hamilton, H.L. A series of normal stages in the development of the chick embryo. 1951. Dev Dyn. 1992 Dec;195(4):231–72.

86. Dai, F., Yusuf, F., Farjah, G.H. & Brand-Saberi, B. RNAi-induced targeted silencing of developmental control genes during chicken embryogenesis. Dev Biol. 2005 Sep 1;285(1):80–90.

87. Scaal, M., Gros, J., Lesbros, C. & Marcelle, C. In ovo electroporation of avian somites. Dev Dyn. 2004 Mar;229(3):643–50.

88. Rehimi, R. et al. Epigenomics-Based Identification of Major Cell Identity Regulators within Heterogeneous Cell Populations. Cell Rep. 2016 Dec 13;17(11):3062–3076.

89. Nieto, H., Patel, D.G. & Wilkinson, M.A. In situ hybridization analysis of chick embryos in whole mount and tissue sections. Methods Cell Biol. 1996;51:219–35.

90. Li, H. & Durbin, R. Fast and accurate short read alignment with Burrows-Wheeler transform. Bioinformatics. 2009 Jul 15;25(14):1754–60.

91. Li, H. et al. The Sequence Alignment/Map format and SAMtools. Bioinformatics. 2009 Aug 15;25(16):2078–9.

92. Broad Institute. (Accessed: Jun 23, 2016). “Picard Tools.” Broad Institute, GitHub repository. http://broadinstitute.github.io/picard/.

93. Ramírez, F., Dündar, F., Diehl, S., Grüning, B.A. & Manke, T. deepTools: a flexible platform for exploring deep-sequencing data. Nucleic Acids Res. 2014 Jul;42(Web Server issue):W187–91.

94. Haeussler, M. et al. The UCSC Genome Browser database: 2019 update. Nucleic Acids Res. 2019 Jan 8;47(D1):D853–D858.

95. Merchant, N. et al. The iPlant Collaborative: Cyberinfrastructure for Enabling Data to Discovery for the Life Sciences. PLoS Biol. 2016 Jan 11;14(1):e1002342.

96. Zhang, Y. et al. Model-based analysis of ChIP-Seq (MACS). Genome Biol. 2008;9(9):R137.

97. Kazachenka, A. et al. Identification, Characterization, and Heritability of Murine Metastable Epialleles: Implications for Non-genetic Inheritance. Cell. 2018;175(5):1259–1271.e13.

98. Quinlan, A.R. BEDTools: The Swiss-Army Tool for Genome Feature Analysis. Curr Protoc Bioinformatics. 2014 Sep 8;47:11.12.1–34.

99. Barrett, T. et al. NCBI GEO: archive for functional genomics data sets--update. Nucleic Acids Res. 2013 Jan;41(Database issue):D991–5.

100. Langmead, B. & Salzberg, S.L. Fast gapped-read alignment with Bowtie 2. Nat Methods. 2012 Mar 4;9(4):357–9.

101. Servant, N. et al. HiC-Pro: an optimized and flexible pipeline for Hi-C data processing. Genome Biol. 2015 Dec 1;16:259.

102. Lareau, C.A. & Aryee, M.J. hichipper: a preprocessing pipeline for calling DNA loops from HiChIP data. Nat Methods. 2018 Feb 28;15(3):155–156.

103. Lareau, C.A. & Aryee, M.J. diffloop: a computational framework for identifying and analyzing differential DNA loops from sequencing data. Bioinformatics. 2018 Feb 15;34(4):672–674.

104. Li, D., Hsu, S., Purushotham, D., Sears, R.L. & Wang, T. WashU Epigenome Browser update 2019. Nucleic Acids Res. 2019 Jul 2;47(W1):W158–W165.

105. Abdennur, N. & Mirny, L.A. Cooler: scalable storage for Hi-C data and other genomically labeled arrays. Bioinformatics. 2020 Jan 1;36(1):311–316.

106. Flyamer, I.M., Illingworth, R.S. & Bickmore, W.A. Coolpup.py: versatile pile-up analysis of Hi-C data. Bioinformatics. 2020 May 1;36(10):2980–2985.

107. Akdemir, K.C. & Chin, L. HiCPlotter integrates genomic data with interaction matrices. Genome Biol. 2015 Sep 21;16(1):198.

108. Zhang, W. et al. The BAF and PRC2 Complex Subunits Dpf2 and Eed Antagonistically Converge on Tbx3 to Control ESC Differentiation. Cell Stem Cell. 2019 Jan 3;24(1):138–152.e8.

109. Tavares, L. et al. RYBP-PRC1 complexes mediate H2A ubiquitylation at polycomb target sites independently of PRC2 and H3K27me3. Cell. 2012 Feb 17;148(4):664–78.

110. Dixon, J.R. et al. Topological domains in mammalian genomes identified by analysis of chromatin interactions. Nature. 2012 Apr 11;485(7398):376–80.

111. Mohammed, H. et al. Single-Cell Landscape of Transcriptional Heterogeneity and Cell Fate Decisions during Mouse Early Gastrulation. Cell Rep. 2017 Aug 1;20(5):1215–1228.

112. Pijuan-Sala, B. et al. A single-cell molecular map of mouse gastrulation and early organogenesis. Nature. 2019 Feb;566(7745):490–495.

## Supplementary References

1. Bleckwehl, T. et al. Enhancer priming by H3K4 methylation safeguards germline competence. bioRxiv. 2020.

2. Zheng, H. et al. Resetting Epigenetic Memory by Reprogramming of Histone Modifications in Mammals. Mol Cell. 2016 Sep 15;63(6):1066–79.

3. Cruz-Molina, S. et al. PRC2 Facilitates the Regulatory Topology Required for Poised Enhancer Function during Pluripotent Stem Cell Differentiation. Cell Stem Cell. 2017 May 4;20(5):689–705.e9.

4. Bejerano, G. et al. Ultraconserved elements in the human genome. Science. 2004 May 28;304(5675):1321–5.

5. Habibi, E. et al. Whole-genome bisulfite sequencing of two distinct interconvertible DNA methylomes of mouse embryonic stem cells. Cell Stem Cell. 2013 Sep 5;13(3):360–9.

6. Zylicz, J.J. et al. Chromatin dynamics and the role of G9a in gene regulation and enhancer silencing during early mouse development. Elife. 2015 Nov 9;4:e09571.

7. Zhang, Y. et al. Dynamic epigenomic landscapes during early lineage specification in mouse embryos. Nat Genet. 2018 Jan;50(1):96–105.

8. Boulard, M., Edwards, J.R. & Bestor, T.H. FBXL10 protects Polycomb-bound genes from hypermethylation. Nat Genet. 2015 May;47(5):479–85.

9. Durinck, S., Spellman, P.T., Birney, E. & Huber W. Mapping identifiers for the integration of genomic datasets with the R/Bioconductor package biomaRt. Nat Protoc. 2009;4(8):1184–91.

